# Integrated single-cell multiomics uncovers foundational regulatory mechanisms of lens development and pathology

**DOI:** 10.1101/2023.07.10.548451

**Authors:** Jared A Tangeman, Sofia M Rebull, Erika Grajales-Esquivel, Jacob M Weaver, Stacy Bendezu-Sayas, Michael L Robinson, Salil A Lachke, Katia Del Rio-Tsonis

## Abstract

Ocular lens development entails epithelial to fiber cell differentiation, defects in which cause congenital cataract. We report the first single-cell multiomic atlas of lens development, leveraging snRNA-seq, snATAC-seq, and CUT&RUN-seq to discover novel mechanisms of cell fate determination and cataract-linked regulatory networks. A comprehensive profile of *cis*- and *trans*-regulatory interactions, including for the cataract-linked transcription factor MAF, is established across a temporal trajectory of fiber cell differentiation. Further, we divulge a conserved epigenetic paradigm of cellular differentiation, defined by progressive loss of H3K27 methylation writer Polycomb repressive complex 2 (PRC2). PRC2 localizes to heterochromatin domains across master-regulator transcription factor gene bodies, suggesting it safeguards epithelial cell fate. Moreover, we demonstrate that FGF hyper-stimulation *in vivo* leads to MAF network activation and the emergence of novel lens cell states. Collectively, these data depict a comprehensive portrait of lens fiber cell differentiation, while defining regulatory effectors of cell identity and cataract formation.

## Introduction

Expression profiling of specific tissues and cell types has been applied for gene discovery in organogenesis^1^. Over the past two decades, genome-level approaches have been applied to study the ocular lens from different species, namely, human, mouse, chicken, amphibian, and fish. These studies involved the use of expression microarrays and high-throughput RNA-sequencing^2–40^. However, the majority of these studies were performed on whole lens tissue, with the exception of a few studies that generated transcriptome data on isolated lens epithelium or fiber cells^9, 22, 33, 35, 41^ or specific regions within the lens^7, 18, 42, 43^. In addition to these omics-level approaches, targeted expression profiling for genes of interest has been performed on individual lens fiber cells in mouse^44, 45^. However, the advent of single cell RNA-sequencing (scRNA-seq) and single nucleus RNA-sequencing (snRNA-seq) now offer new opportunities to examine global gene expression from single cells or nuclei at an unprecedented throughput and resolution^46–50^.

Even so, the application of these new approaches to study the developing lens have been limited^51^. Indeed, except for recent studies in zebrafish^52^ and mouse^53^, there are no other reports of scRNA-seq or snRNA-seq on the developing lens. The zebrafish study, while broadly informative, performed scRNA-seq on the whole embryo, thus limiting the number of cells and depth of the analysis related to the lens^52^. In the mouse study, scRNA-seq was performed on the lens placode at the lens-induction stage^53^. Thus, while informative for examining gene expression profiles associated with early lens development, the mentioned study does not inform on gene expression dynamics in different lens epithelial cell populations, or on the differentiation of cells in the transition zone into fiber cells and their sub-populations. In contrast, while the chicken has been used as a model for lens development for several decades^54–56^, studies on defining the transcriptome of chick lenses have been limited^42, 43^. The two transcriptomics studies on the chick lens have examined the embryonic lens at the resolution of distinct regions within the tissue^42, 43^, but not at the single cell level. In these studies, bulk RNA-seq was performed on micro-dissected lens regions representing the centrally- and equatorially-located epithelia, and the cortically- and centrally-located fiber cells^42, 43^. In addition to RNA-seq, the latter study also described global chromatin accessibility profiles related to these distinct lens regions, thus allowing correlation between the state of the chromatin and RNA abundance in a coarse spatial manner across the tissue^43^. Other studies performed in the mouse similarly focused on defining the correlation between chromatin state and gene expression in either whole lenses^57^ or the isolated epithelium and fiber cells^58, 59^. While helpful, advanced technologies now allow new opportunities to investigate developing tissues and their underlying molecular networks at single-cell resolution.

To advance knowledge on the intersection of chromatin state dynamics and gene expression in the lens at single-cell level, we present an integrated, multiomic approach that encompasses a global profile of gene expression and chromatin accessibility associated with the chicken epithelial to fiber cell differentiation program (Fig. 1a). We generate and analyze multiple omics modalities and layer functional approaches to define the molecular cascades underlying lens development and their significance to lens pathology. Specifically, we capture epithelial cell state dynamics and establish an inferred temporal map of the epithelial to fiber cell differentiation trajectory. This approach not only confirms signal established effectors active in lens development, such as, FGF, WNT, GLI, and NOTCH, but further informs on their precise spatiotemporal dynamics in lens development. The multimodal viewpoint allows us to construct a comprehensive atlas of the *cis*- and *trans*-regulatory networks that hallmark epithelial cell dynamics and fiber cell differentiation. To this end, we identify established lens transcription factors (TFs), such as PAX6, FOXE3, MAF, and SOX-family proteins, among others, and predict new candidates for future study. We leverage these observations to deduce global binding events and downstream regulatory targets for the cataract-linked transcription factor MAF, a known effector of lens fiber cell differentiation. Additionally, this integrated analysis uncovered a novel, conserved epigenetic paradigm of fiber cell differentiation, defined by loss of the H3K27 methylation writer Polycomb Repressive Complex 2 (PRC2) and the simultaneous acquisition of the PRC2-modifying gene *JARID2*^60–62^. We profile the global DNA-binding activity of PRC2 and the distribution of histone modifications to uncover a potential new epigenetic mechanism for safeguarding early lens cell identity. Finally, using FGF2 beads, we examine the functional effects of FGF hyper-stimulation on lens cell identity *in vivo* and at single-cell resolution. Together, these rich datasets confirm established pathways and known cataract-linked genes in the lens, and through independent validation and functional assays, we identify a cohort of new proteins that play a key role in lens development, homeostasis, and pathology.

**Fig. 1.**
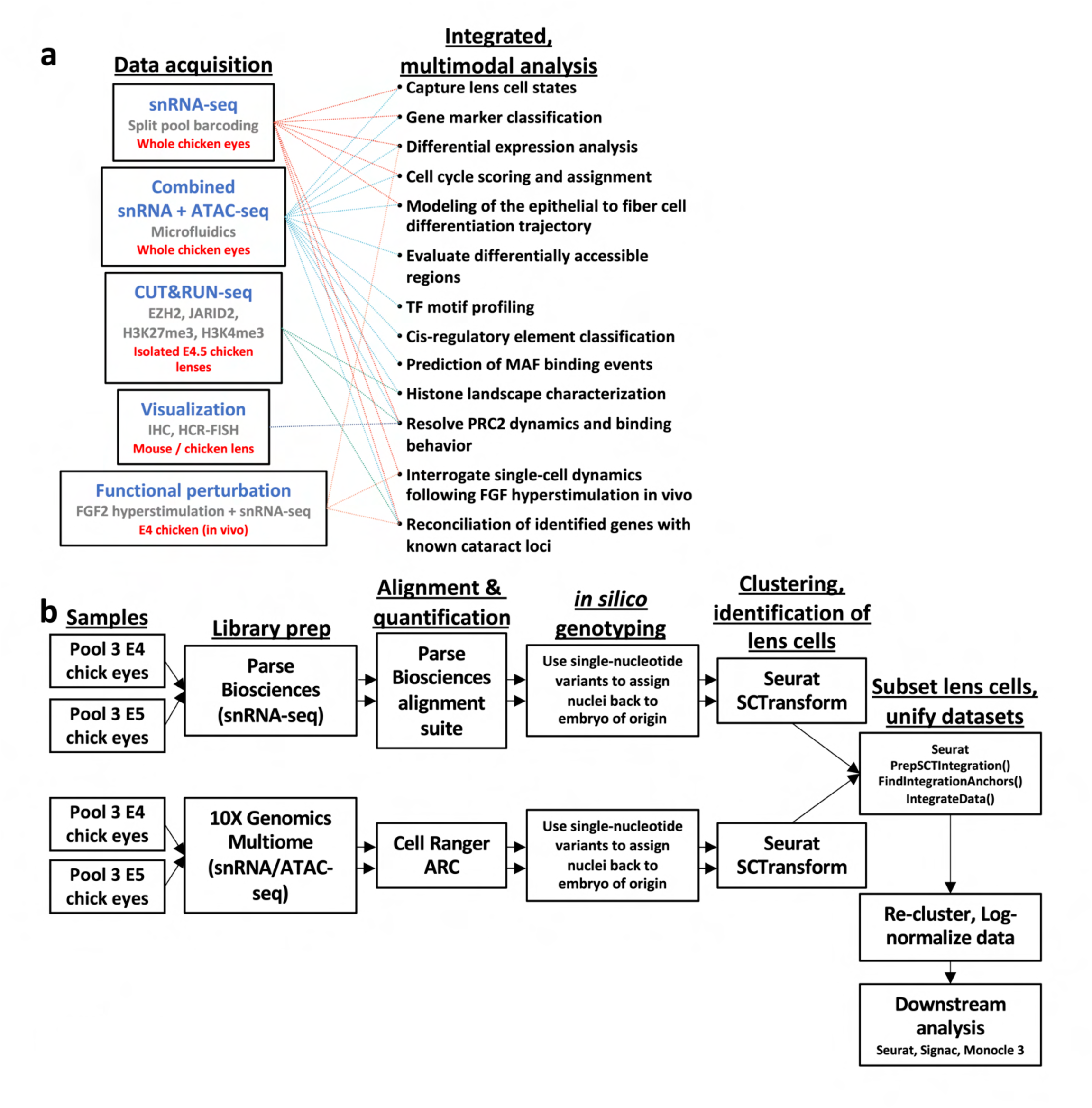
Experimental overview. (**a**): Schematic outlines the major methodology and analysis employed to capture an integrated, multimodal portrait of lens development. IHC = immunohistochemistry; HCR-FISH = Hybridization Chain Reaction-Fluorescent *In-Situ* Hybridization; TF = Transcription Factor. (**b**): Workflow summarizes major steps in the generation of single-nuclei libraries, as well as the initial processing of snRNA-seq data and identification of lens cells.

## Results

### Single-nuclei profiling identifies distinct cell populations in the embryonic chicken lens

We sought to examine the global regulatory profile of the embryonic chicken lens on a multimodal, single-nuclei level (Fig. 1a). We established a working protocol to process and identify lens nuclei by performing high-throughput RNA-sequencing (snRNA-seq) and ATAC-sequencing (ATAC-seq) on the whole chicken eye, which can inform on the transcriptome and chromatin landscape (Fig. 1b). Briefly, whole eyes were isolated from chickens at embryonic (E) day 4 and 5, processed using an in-house nuclei isolation protocol for library preparation, and were subjected to either snRNA-seq or combined snRNA-seq and ATAC-seq. From the snRNA-seq profiles, which comprised of a total 20,066 nuclei, 643 nuclei were identified by a UMAP feature plot to exhibit characteristic lens gene expression, evinced by progressive abundance of *ASL1*, which encodes delta-crystallin, an established marker for the chicken lens (Fig. 2a). Based on re-clustering of this isolated lens population, this population could be separated into three major cell states (Fig. 2b). Next, we examined these subclusters by applying an approach termed “cell cycle scoring” that – based on the abundance of cell cycle-related transcripts – assigns a specific cell cycle stage to each nucleus (Fig. 2b). This analysis, as well as the progressive abundance of *ASL1*, led to the recognition of the lens subclusters as of epithelial, intermediate, and fiber cell origin (Fig. 2b). Each of these subclusters was comprised of nuclei derived from each of two distinct library preparation technologies, as well as from distinct biological replicates, and both sexes (Supplementary Fig. 1). In agreement, the progressive reduction or gain of specific genes associated with epithelial to fiber transition were observed in these subclusters. For example, *TFAP2A*, *DMRTA2*, and *FOXE3* were abundant in the epithelial subcluster and were progressively reduced in the intermediate and fiber subclusters (Fig. 2c). Conversely, *MIP*, *BFSP1*, and *CRYBA4* were largely absent in the epithelial subcluster and were progressively abundant in the intermediate and/or fiber subclusters (Fig. 2c). In total, 1,417 unique genes were found to be selectively enriched in any of the 3 identified subclusters (Supplementary File S1). Further, the recognition of these epithelial, intermediate and fiber subclusters were independently supported by pathway enrichment analysis. Indeed, functional pathways associated with genes enriched in the epithelial subcluster included “mitotic cell cycle process” and “morphogenesis of an epithelium” (Fig. 2d; Supplementary File S2). Terms associated with the intermediate subcluster-enriched genes included “establishment or maintenance of cell polarity”, while those associated with the fiber subcluster included “lens fiber cells” (Fig. 2d). Together, these analyses demonstrate that the snRNA-seq assay reliably identifies lens cells and classifies them into populations reflecting epithelial, intermediate, and fiber cell biology.

**Fig. 2.**
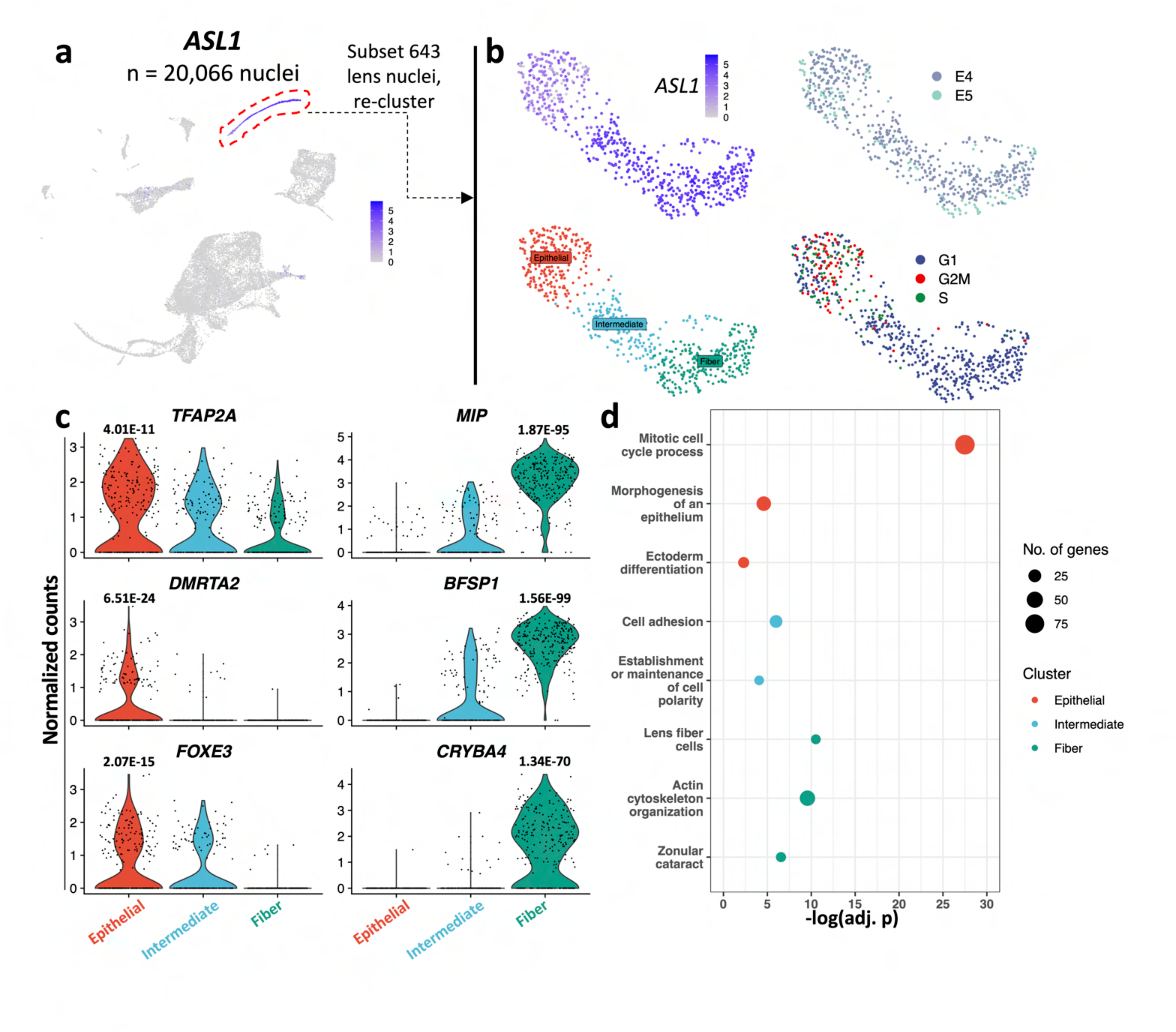
Single-nucleus profiling of the chick lens. (**a**) UMAP feature plot displays *ASL1* transcript abundance, in log-normalized gene counts, in nuclei collected from the whole eye of developing chicken embryos. (**b**) Lens nuclei were subset and re-clustered, revealing 3 major cell states. Cells are colored by *ASL1* RNA abundance, cluster, stage, or cell cycle phase. (**c**) Violin plots display the distribution of log-normalized RNA abundance for genes related to epithelial (on the left) or fiber cell identity (on the right). Adjusted p-values above violin plots represent gene marker enrichment, calculated via Wilcoxon Rank Sum test applied to each cluster relative to other two cell populations. (**d**): Pathway enrichment summarizes major biological functions associated with marker genes for each cluster.

### Derivation of the lens epithelial to fiber cell differentiation trajectory by snRNA-seq

While clustering analysis was able to coarsely define lens cell populations, we postulated that the biology of these cells may be more aptly considered as a continuum of expression states, rather than as discrete populations. To this end, we employed pseudotime analysis to establish a continuous trajectory of epithelial to fiber cell differentiation. Pseudotime analysis is an analytic framework that assigns a measure of how far a cell has moved across a biological process. Using this approach, cells were ordered by their global expression profiles over an inferred progression of epithelial to fiber cell identity (Fig. 3a). Viewed from this perspective, the established trajectory concurred with previously accepted expression patterns reflecting epithelial through fiber cell differentiation, in turn, allowing us to infer temporal aspects of gene regulation across lens cell states. For example, *EFNA5 (ephrin-5A)* is abundant early and gets progressively reduced in this inferred epithelial to fiber trajectory (Fig. 3b), which is consistent with its established expression in the lens^63^. Similarly, other genes, such as *CRIM1*, *TGFB2*, *TRPM1*, previously known to show this pattern^63, 64^, were identified in this analysis (Fig. 3b). Further, in agreement with their established expression patterns^33, 42, 63, 65, 66^, *BIN1*, *BNIP3L*, *CAPRIN2*, *EPHA2*, *GJA3*, *GJA8*, and *RBM24* exhibited progressive abundance across the epithelial to fiber continuum (Fig. 3b). Consistent with the known literature^67^, *JAG1* is found to exhibit transient high expression in the epithelial to fiber continuum. It should be noted that the total number of genes correlating with epithelial and fiber states were significantly higher compared to the number correlating with the intermediate state (Supplementary File S1). This analysis also identified novel candidates with spatially dynamic expression patterns for future studies in the lens. Among others, these included the solute carrier protein *SLC7A2*, the transmembrane protein *TMEM178B*, the predicted Calcium ion binding protein *CLSTN2*, the ephrin ligand *EFNB2*, the fibroblast growth factor receptor like protein *FGFRL1*, and the cytoskeletal protein spectrin *SPTBN1*. Taken together, the captured trajectory encapsulates numerous previously reported features of the epithelial to fiber cell differentiation program, while also broadly informing on novel spatiotemporal dynamics of RNA regulation.

**Fig. 3.**
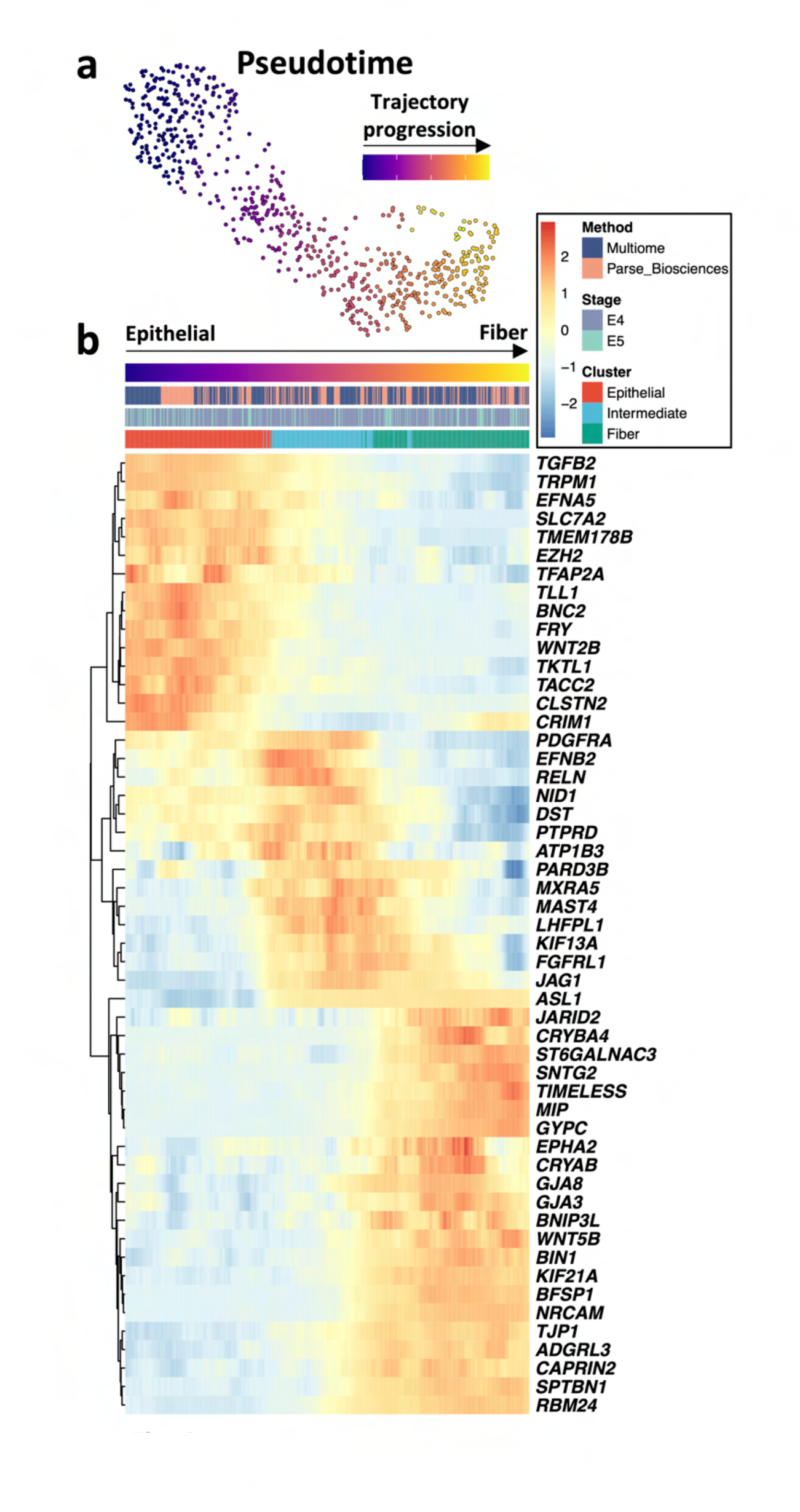
An inferred temporal map of epithelial to fiber cell differentiation. (**a**): UMAP displays lens nuclei color-coded by pseudotime value. (**b**): Heatmap displays the expression of select gene markers for epithelial, intermediate, or fiber cell identity along the inferred differentiation trajectory.

### snATAC-seq captures chromatin configuration during epithelial to fiber cell differentiation

Next, we sought to define the chromatin changes associated with the epithelial through fiber differentiation program. Toward this goal, we took a multiomics approach that facilitates the simultaneous capture of RNA and accessible chromatin from the same nucleus. Using this approach, in addition to the RNA profiles described above, we also captured the ATAC profiles of 386 nuclei across the lens population. These combined RNA and ATAC profiles are summarized by UMAP representation (Fig. 4a). Notably, the combined modality UMAP preserves the spatial proximity of cells within the subclusters previously derived from the RNA-only profiles. In addition, all nuclei clusters displayed expected enrichment patterns proximal to transcription start sites (TSS) and nucleosomal periodicity across the distribution of aligned fragment lengths (Supplementary Fig. 2). Differential accessibility testing of ATAC-seq peak regions identified 963 peak regions that exhibit a significant change in chromatin accessibility during fiber cell differentiation. This analysis informed on the chromatin accessibility of differentiating lens cells, which broadly correlated with epithelial and fiber states (Fig. 4b).

**Fig. 4.**
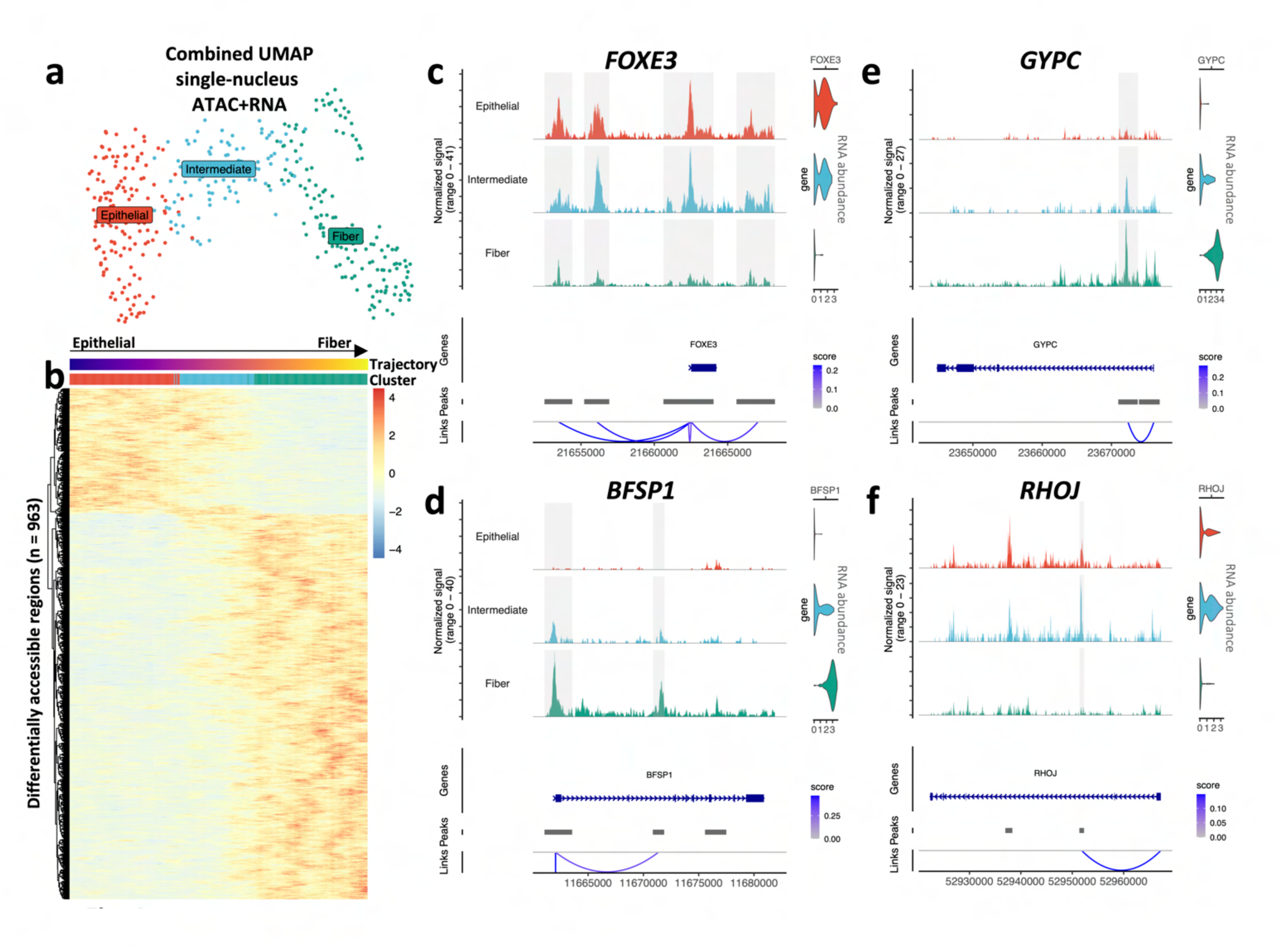
Cis-regulatory modules of the lens differentiation program. (**a**): UMAP generated from weighted nearest neighbor analysis of RNA and ATAC profiles of lens nuclei. (**b**): Row-normalized ATAC signal within differentially accessible regions are displayed across the pseudotime trajectory. (**c-f**): Genome browsers display accessibility signal plotted across the loci of marker genes for each subcluster. The structure of the gene body is indicated by blue bars, with thick regions corresponding to exons, and arrows oriented toward direction of coding sequence. Link tracks on bottom display predicted looping interactions between peak regions and transcription start sites. The links are colored by the link score, which corresponds to the strength of the predicted association between accessibility and expression. Accessible peak regions linked to transcription are highlighted. Normalized RNA abundance for each gene is displayed in violin plots on right. Numbers along the bottom of each panel are genomic coordinates.

Given these observed changes in chromatin organization, we next sought to leverage our multimodal approach to correlate changes in peak accessibility with changes in the expression of nearby genes. Based on this logic, we used normalized expression and chromatin accessibility to predict interactions between peak regions and TSS, limiting these interactions to flanking the TSS within 1 million base-pairs. This analysis identified potential *cis*-regulatory elements (CREs) that correlate with the dynamic expression pattern for individual genes. For example, three distal CREs and a peak spanning the coding region were identified for *FOXE3*, and the accessibility of these potential regulatory regions correlated positively with *FOXE3* expression dynamics (*i.e.*, high expression and accessibility in the epithelial state that are both progressively reduced in the fiber state) (Fig. 4c). Similarly, an expression-linked CRE and promoter region peak were also identified for the fiber cell-expressed gene *BFSP1* (Fig. 4d). Interestingly, this analysis also identified numerous fiber-expressed genes as promising candidates for further study in the lens (*e.g.*, the glycophorin-C encoding gene *GYPC*) (Fig. 4e). Importantly, this approach was sensitive enough to identify genes whose dynamic expression spiked ephemerally in the intermediate state, such as *RHOJ*, which encodes a Ras homolog family member GTP-binding protein, as well as a transiently accessible CRE predicted to drive the transcription of *RHOJ* (Fig. 4f). The *cis*-regulatory profile of additional known – as well as novel – lens regulators are displayed, including *ASL1*, *CRYAB*, *CRYBA4*, *PAX6*, *TKTL1*, and *YBX3* (Supplementary Fig. S3). In a similar fashion, we report the expression profile and chromatin landscape of genes that serve as effectors of pathways with known roles in mediating lens fate, including FGF, BMP, SHH, and NOTCH signaling, as well as numerous mediators of intracellular cascades (Supplementary Fig. S4).

Additionally, we postulated that this analysis could inform on the dynamic expression pattern of the genes encoding TFs in lens development. We used established repositories^68, 69^ and manual curation for analyzing mouse and human TFs that are expressed in the lens. This approach identified 80 TF-encoding genes with a dynamic expression pattern in lens epithelial through fiber cell states (Fig. 5a). Importantly, this analysis identified key lens TFs (*e.g.*, *FOXE3*, *HSF4*, *MAF*, *MAB21L1*, *MEIS2*, *PAX6*, *PROX1*, *SOX1*, and *SOX2*) and could capture their established expression patterns in epithelial through fiber differentiation (Fig. 5a)^70^. For example, *PAX6* levels are high early and are reduced in this inferred epithelial to fiber trajectory (Fig. 5a), consistent with its known expression pattern in the lens^71^. Next, we sought to relate the expression dynamics of the genes encoding lens TFs to the chromatin signature observed at their cognate DNA-binding motifs throughout the genome. Toward this goal, we used the JASPAR database^72^ to procure known vertebrate TF motifs and searched for their overrepresentation within accessible genomic regions across specific lens states. This approach identified a global reduction in accessibility in the genomic regions that contain a FOX-family motif in epithelial to fiber differentiation (Fig. 5b). This correlates with expression of *FOXE3*, which is progressively reduced as cells transition from epithelial to fiber states. This analysis also identified shifts in motif accessibility for known TFs (*e.g.*, E2F3)^73^ as well as for contextually novel TFs in the lens (*e.g.*, the E2F-binding protein TFDP1)^74^ that follow a similar pattern throughout epithelial to fiber transition. Similarly, this analysis identified a global elevation in accessibility within genomic regions that contain MAF and SOX13 motifs as cells transition from epithelial to fiber states, which agrees with the observed gains in RNA abundance observed for both of these factors (Fig. 5c). Interestingly, this analysis points to a previously unappreciated role for SOX2 in fiber differentiation, which exhibits elevated RNA abundance and motif accessibility in the fiber cell population (Fig. 5c).

**Fig. 5.**
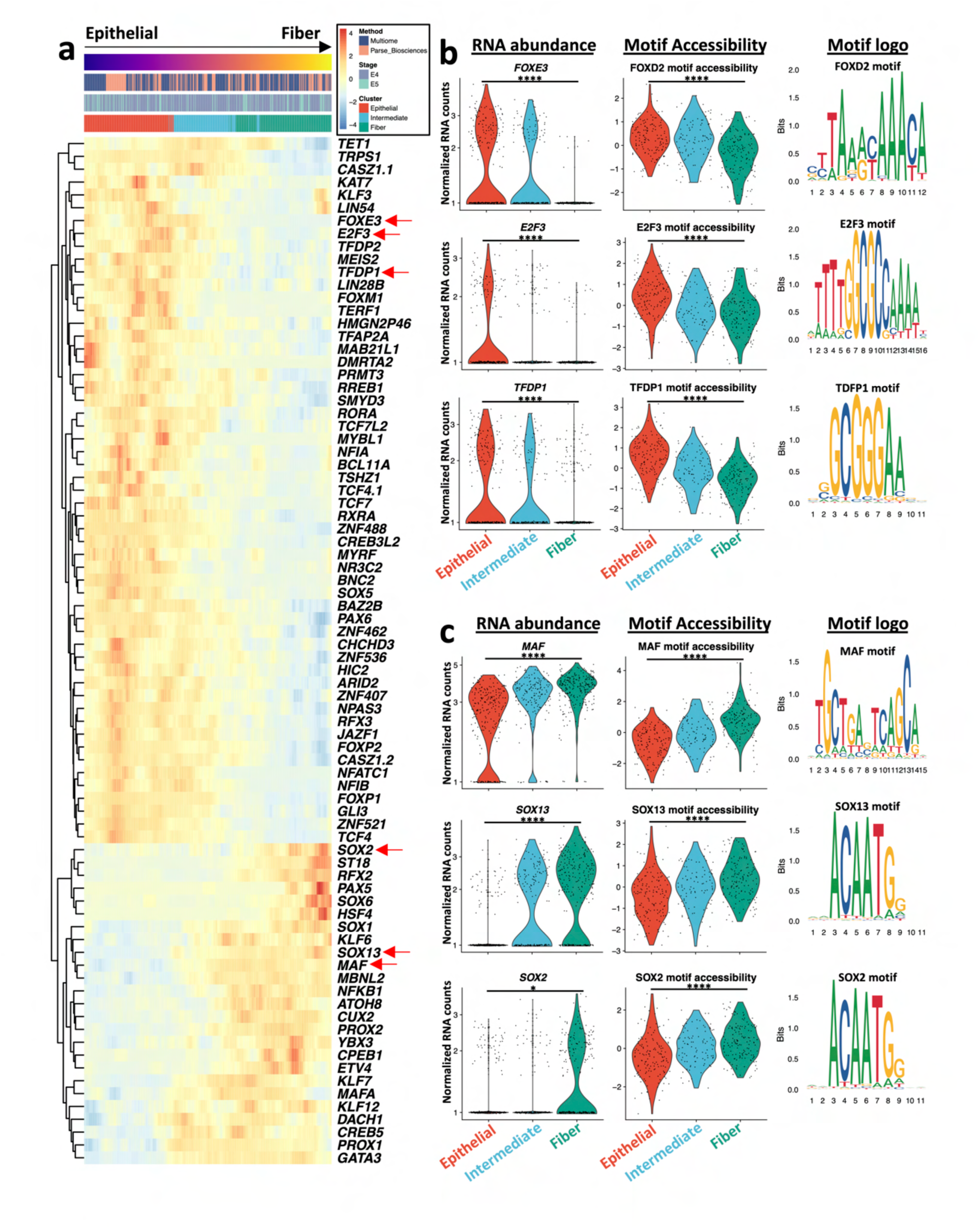
Global chromatin footprints imparted by the trans-effectors of epithelial vs. fiber cell identity. (**a**): Trans-effectors of the lens differentiation program. Heatmap displays pseudo-temporal regulatory expression patterns of a manually curated list of transcription factor-encoding genes regulated during lens fiber cell differentiation. (**b**): Left violin plots display log-normalized RNA abundance for genes down-regulated in fiber cells within each cell population. Right violin plots display the chromVAR enrichment scores for the cognate motif within accessible peak regions. Motifs are derived from the JASPAR database. (**c**): Same as B but displaying genes up-regulated in fiber cells. Asterisk (*) denotes adjusted p-value < 0.05; (****) denotes adjusted p < 0.00005; calculated via Wilcoxon Rank Sum test comparing Epithelial vs. Fiber clusters.

### Derivation of regulatory networks for key lens transcription factors associated with human ocular defects

Next, we sought to apply the chromatin accessibility information to predict regulatory relationships with specific transcription factors in the lens. As a proof of principle, we focused on the bZIP family protein MAF (c-Maf), mutations or deficiency of which cause cataract and other ocular defects in humans^75–77^ and in animal models^78–80^. We took the following approach toward this goal: (1) identify all peak regions correlated with a nearby gene expression change event, (2) identify the subset that contain Maf-binding motif(s), (3) identify the subset that exhibit a predicted activating relationship (*i.e.*, upregulation of nearby gene expression), and (4) retain only those peaks that also have a moderate positive correlation between accessibility and *MAF* RNA abundance. Cells across the epithelial through fiber states exhibit a progressive increase in *MAF* expression, which correlate with increased accessibility of specific genomic regions containing a MAF binding motif (Fig. 6a). We searched for linked genes within 1 million base-pairs of these MAF motif-containing genomic regions; of these, 114 genes met all of the defined selection criteria, suggestive of an activating relationship with MAF binding (Fig. 6b). These candidates included several key cataract/lens defect-linked genes as putative MAF targets in the lens, including *BFSP2*, *CRYBB1*, *GATA3*, and *TDRD7* (Fig. 6c), among others (Supplementary Fig. S5). In addition to these candidates, other genes were identified with important roles in lens biology (*e.g.*, *ASL1*, *CAP2*, *CAPRIN2*, *CRYBA1*, *CRYBA4*, *CRYBB2*, *CRYBB3*, *JAG1*, *MAFA*, *PROX1*, *RBM24*, *SOX13*). Notably, using this approach, MAF itself is identified among the target genes, which is indicative of auto-regulatory circuitry. Critically, MAF has been demonstrated to bind proximal to its own locus and potentiate its expression in the developing mouse lens, as well as to drive the expression of *Crybb1*^70, 81, 82^. Thus, our predictive model identifies accepted *bona fide* MAF targets, while expanding the repertoire of candidate MAF targets that are active in the lens for future exploration. Altogether, this represents a multiomics-based heuristic approach for predicting MAF regulatory outputs in lens development and pathology, and a similar method could likely be extended to other key TFs.

**Fig. 6.**
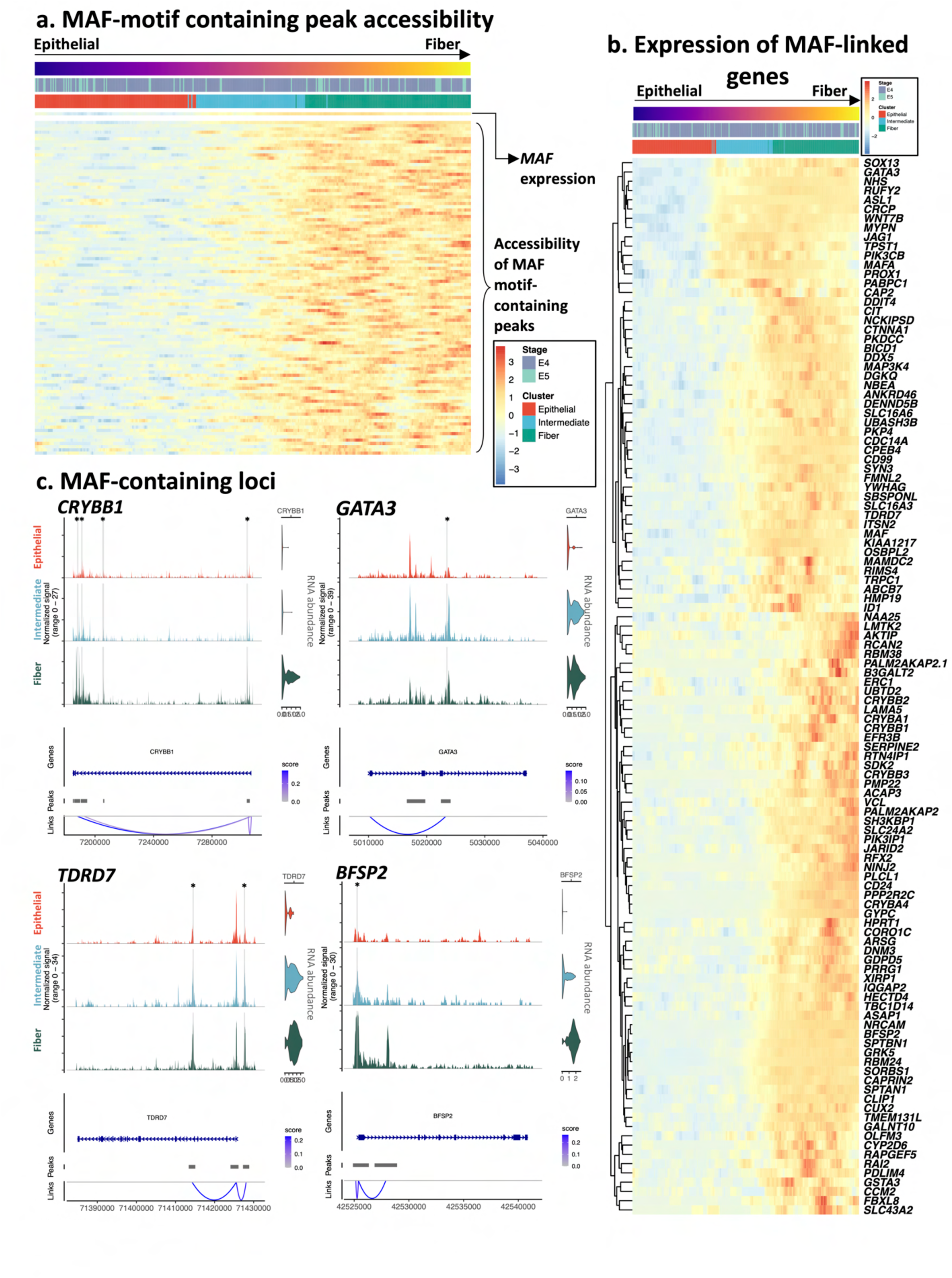
Derivation of the MAF regulatory network. (**a**): Top row of row-normalized heatmap displays RNA abundance of *MAF* RNA transcripts across the pseudotime trajectory. Rows below display accessibility of MAF motif-containing loci linked to nearby changes in transcription. (**b**): The RNA abundances of MAF-linked genes are shown. (**c**): The loci of genes containing MAF-linked peaks are displayed, with MAF motif-containing peaks highlighted and marked by an asterisk. Accessibility signal is plotted across the loci. The structure of the gene body is indicated by blue bars, with thick regions corresponding to exons, and arrows oriented toward direction of coding sequence. Link tracks on bottom display predicted looping interactions between peak regions and transcription start sites. The links are colored by the link score, which corresponds to the strength of the predicted association between accessibility and expression. Normalized RNA abundance for each gene is displayed in violin plots on right. Numbers along the bottom of each subpanel are genomic coordinates.

### Identification of a novel, conserved epigenetic program associated with epithelial to fiber cell differentiation

Next, we sought to examine the omics datasets to identify potential novel regulatory events in lens differentiation. Therefore, we focused on genes that exhibit variable expression in the lens epithelial to fiber pseudotime trajectory, which is marked by progressive abundance of *ASL1* (Fig. 7a). Notably, several components of the PRC2 complex, namely *JARID2*, *EZH2*, *EED* and *SUZ12*, were identified among the candidates that had significant variation in expression as epithelial cells transitioned into fiber cell states (Fig. 7a; Supplementary File S3). Of these factors, the three core members of the PRC2 complex, namely *EZH2*, *EED*, and *SUZ12*, exhibit a progressive loss of abundance as cells transition from epithelial to fiber states. The PRC2 complex is a writer of the repressive histone modification H3K27me3 and is recognized as key regulator of gene silencing during development^60, 61^. Conversely, *JARID2* – which encodes a PRC2 accessory protein that can modify the function of this complex – is expressed in all the lens populations but exhibits a progressive elevation of abundance associated with differentiation. This pattern of *EZH2* and *JARID2* mRNA expression in the E4 chicken lens was independently validated by hybridization chain reaction FISH (Fig. 7b). Further, this pattern was conserved between chicken and mouse, as independently validated by examination of previously described transcriptomic data from isolated mouse lens epithelial and fiber cells^33^.

**Fig. 7.**
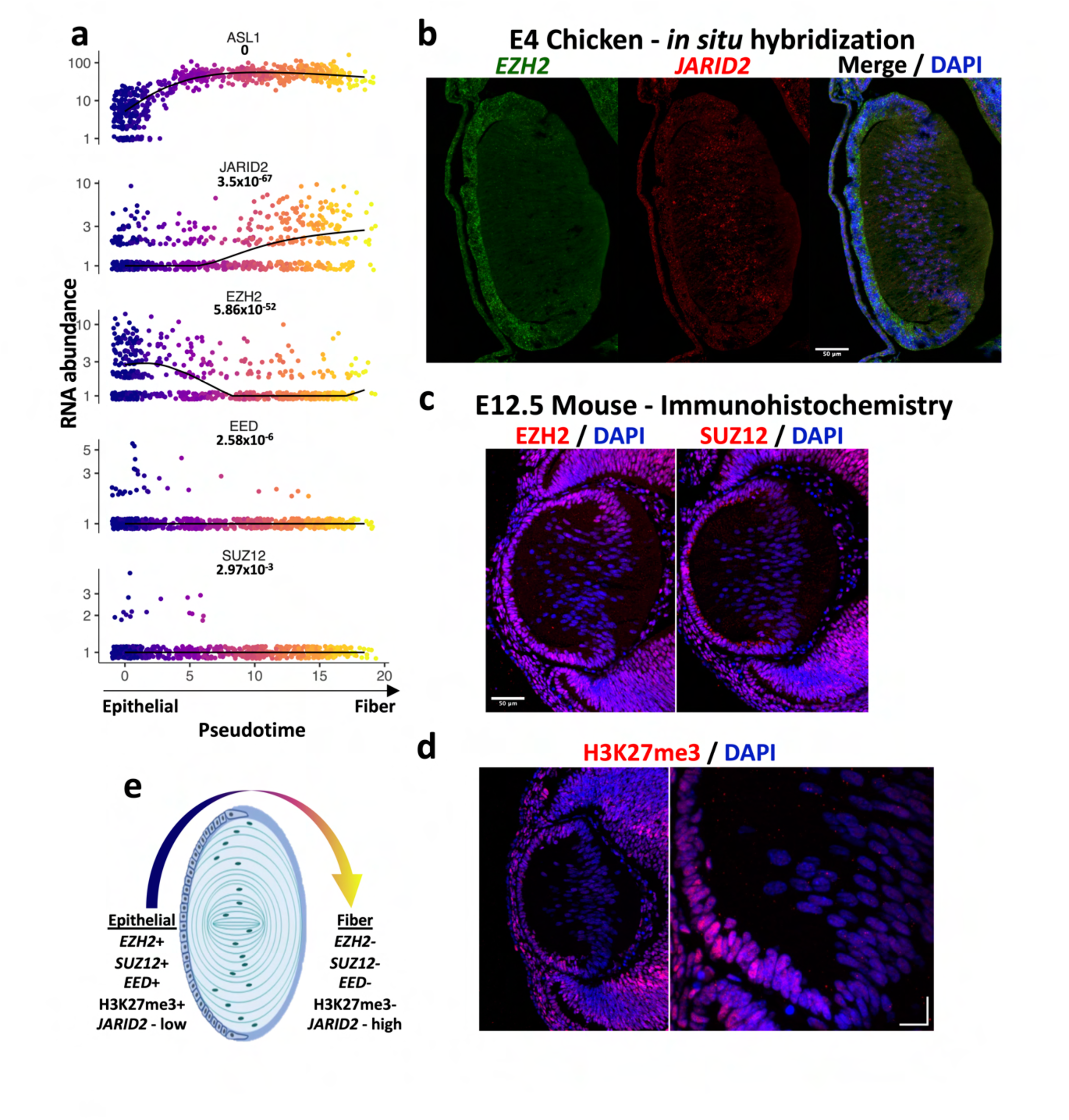
PRC2 dynamics constitute an epigenetic program of fiber cell differentiation. (**a**): The expression of genes encoding PRC2 complex subunits are altered throughout the chick lens fiber cell differentiation program. Nuclei are ordered by pseudotime on the x-axis, and the black line represents the expression trend. Adjusted p-values displayed calculated via Moran’s I test using principal graph. (**b**): FISH-HCR is used to visualize changes in localization of *EZH2* and *JARID2* transcripts in E4 (HH23-24) chick embryos. Scale bar is 50 µm. (**c**): Immunohistochemistry performed on E12 mouse sections show loss of PRC2 members during fiber cell differentiation. Scale bar is 50 µm. (**d**): H3K27me3 is lost from differentiating mouse fiber cells. Scale bar from C applies to left image. Right image scale bar is 15 µm. (**e**): Schematic summarizes changes in PRC2 complex members.

On the protein level, the abundance of EZH2 in E12.5 mouse lens epithelial cells was independently validated by immunofluorescence staining, both of which showed a clear reduction in fiber cells (Fig. 7c). Notably, reductions in SUZ12 abundance were also observed via immunofluorescence staining, although variability was observed amongst biological replicates of the E12.5 mouse lens (Fig. 7c; Supplementary Fig. 6a). However, staining of the E14 mouse lens presented a marked reduction in SUZ12 (Supplementary Fig. 6b), suggesting that this phenotype becomes progressively salient during this window of development. Interestingly, we did not find evidence of differential regulation of *EZH1*, an *EZH2* paralog, in our chick snRNA-seq dataset (Supplementary Fig. 6c). Visualization of EZH1 protein in the developing mouse lens showed EZH1 localized to extra-nuclear deposits along the apical surface of the lens epithelium, and these deposits appeared to redistribute toward the anterior lens as fiber differentiation proceeded (Supplementary Fig. 6d,e). Expectedly, we observe positive H3K27me3 immunofluorescence signal in mouse lens epithelial cells, which was markedly absent from fiber cells (Fig. 7d; Supplementary File S4). Taken together, the overlap of EZH2, SUZ12, and H3K27me3 signals suggests that the PRC2 complex actively functions in rendering spatiotemporal epigenetic control over chromatin structure in lens epithelial cells, an activity which is abrogated from fiber cells. These data suggest a key function for PRC2 complex proteins in cell fate control in lens development (Fig. 7e).

### Genome-wide profiling of PRC2 and histone modifications in the developing lens via CUT&RUN

We next sought to identify the genomic regions where the PRC2 subunits EZH2 and JARID2 are localized in the lens and correlate these with H3K27me3 modification. Toward this goal, we performed CUT&RUN analysis of E4.5 chicken lens. CUT&RUN involves usage of an antibody-guided nuclease to identify global DNA-binding regions for specific proteins of interest^83^. CUT&RUN analysis of embryonic chicken lens showed robust colocalization of signals for EZH2, JARID2, and H3K27me3 (which marks repressed transcription) across their defined peak regions, which were largely devoid of H3K4me3 (which marks active transcription) and the IgG negative control (Fig. 8a, Supplementary Fig. 7a-c). These data suggest that PCR2 is active in facilitating H3K27me3 modifications in lens development. Interestingly, the EZH2/JARID2 co-localized peaks were predominantly identified proximal to genic regions (94.7%) compared to distal intergenic regions (5.3%), implicating a direct function in mediating transcriptional output (Fig. 8b; Supplementary File S5). Within the genic regions, these peaks were most commonly identified in/near the promoter regions (∼75%) (Fig. 8b). Comparison of the top 25 EZH2 or JARID2 bound genes exhibited a 60% overlap (Fig. 8c). Strikingly, almost the entirety of these top bound candidates encoded master-effector transcription factors, transcriptional regulators, or DNA-binding factors involved in cell fate determination, as indicated by their established function in the literature. Indeed, GO analysis of the top 200 PRC2-bound targets demonstrate that a vast majority of these genes are involved in transcription regulation, as indicated by the enrichment of the term “DNA-binding TF activity”, and a high number are also involved in cell fate control, as indicated by the enrichment of the terms “Cell fate commitment” and “Neuron fate commitment” (Fig. 8d; Supplementary File S2).

**Fig. 8.**
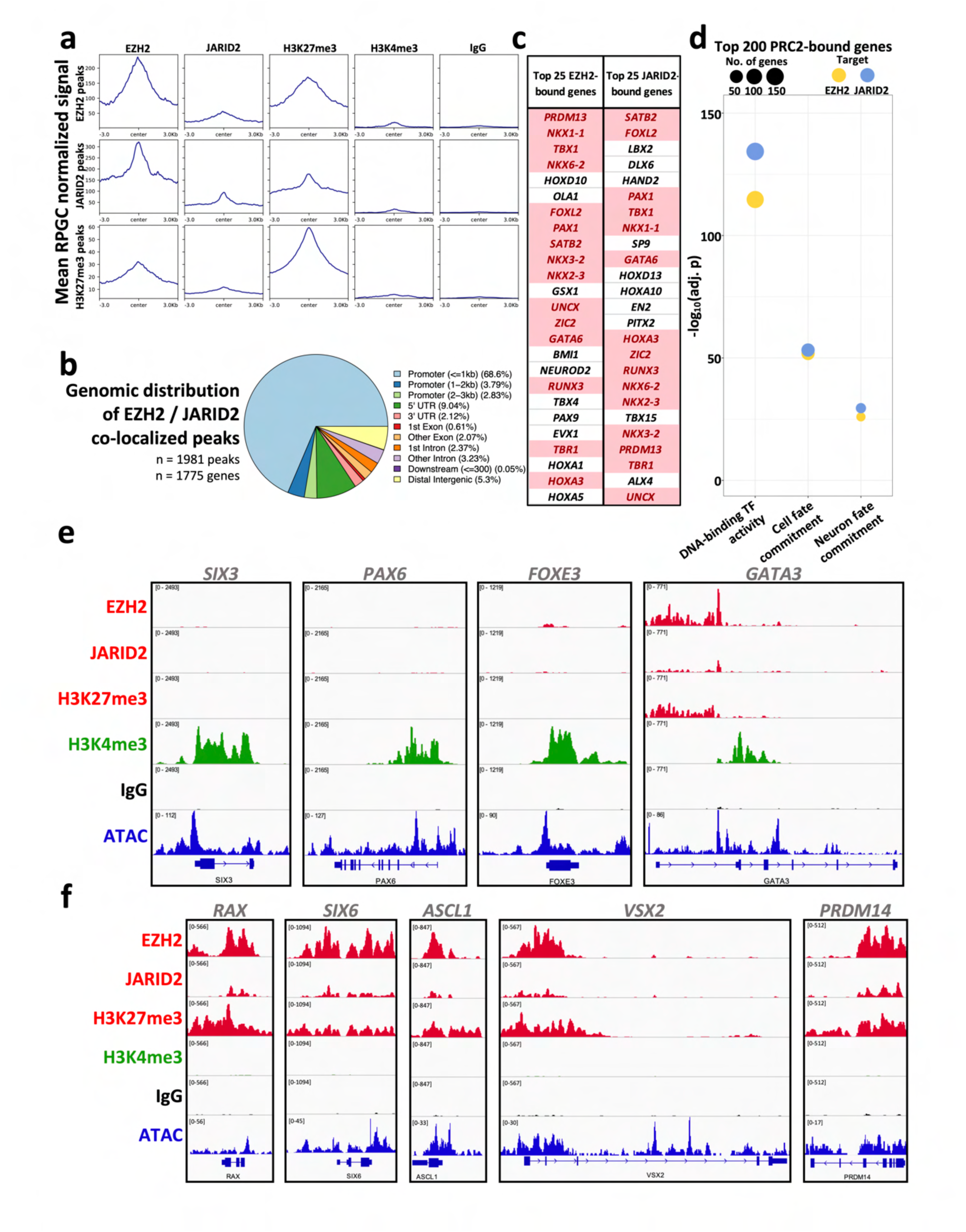
Localization of PRC2 members and histone modifications in the E4.5 chicken lens. CUT&RUN-seq was performed using antibodies against EZH2, JARID2, H3K27me3, H3K4me3, and IgG control. (**a**): Average signal was plotted for the top 1000 peaks identified for EZH2, JARID2, and H3K27me3, revealing a high degree of co-localization between the targets. (**b**): The genomic distribution is displayed for loci co-bound by EZH2 and JARID2, showing predominantly promoter and genic localization. (**c**): Top 25 genes displaying the highest signal for EZH2 and JARID2 are displayed. Duplicated values highlighted in red. (**d**): Bubble chart summarizes pathway enrichment results performed on top 200 genes bound by EZH2 and JARID2. (**e**): Genome browsers display RPGC-normalized (reads per genomic content-normalized) signal for CUT&RUN targets across select loci, as well as aggregate ATAC signal from the snATAC-seq dataset. (**f**): Genome browsers display RPGC-normalized CUT&RUN signal across PRC2-marked loci. Signal ranges are equally scaled for CUT&RUN targets and displayed in top left of each track.

Interestingly, the loci encoding key lens TFs that are known to function in epithelial cells, namely, *PAX6*, *SIX3* and *FOXE3*, did not exhibit enrichment of EZH2, JARID2, or H3K27me3 signal, but did exhibit robust signal for H3K4me3, which is consistent with their active transcription and abundant mRNA levels in the epithelial cell population (Fig. 8e). Similarly, we did observe a small, but notable, number of marker genes associated with subpopulations of our snRNA-seq dataset that were bound by PRC2 (Supplementary Fig. 7d). In this regard, localization patterns between PRC2 and the histone landscape pointed toward sophisticated control over the activation of specific TFs in the larger lens gene regulatory network. For example, *GATA3* exhibits PRC2 peaks while also exhibiting features of open chromatin and transcription activation (Fig. 8e). The totality of our dry and wet lab observations [*e.g.*, (1) PRC2 is bound to the *GATA3* locus, (2) fiber cells exhibit loss of PRC2 and H3K27me3 signals, (3) there is progressive transcriptional activation of *GATA3* in epithelial to fiber differentiation, as indicated by histone modifications (marked by H3K4me3 peaks), (4) there are concomitant changes to the chromatin architecture as demonstrated by ATAC-seq, (5) there are putative MAF-binding sites that we identified within the *GATA3* regulatory locus (Fig. 6c), and (6) progressive abundance of *GATA3* mRNA is observed in the epithelial to fiber snRNA trajectory (Fig. 5a, 6b)], serve to define a complex orchestrated sequence of events that can be reconstructed by this multiomics approach (Supplementary Fig. S8). Conversely, a cohort of non-lens cell expressed TFs, generally involved in cell fate determination of non-lens ocular cell types, exhibited robust localization of EZH2, JARID2, and H3K27me3, but not H3K4me3, demonstrating the PRC2-mediated maintenance of heterochromatin domains that facilitate their transcriptional repression in the lens (Fig. 8f). For example, eye field transcription factors *RAX* and *SIX6*, as well as developmental regulators of neural retina fates, such as *ASCL1*, *VSX2*, and *PRDM14*, all exhibited extensive domains of PRC2 and H3K27me3 across their respective transcript start sites (Fig. 8f). Together, this data suggests that PRC2 function is actively involved in the control of differentiation and the suppression of alternate cell fates in the lens.

### snRNA-seq analysis of the impact of FGF on lens cell fate

We next sought to gain *in vivo*, mechanistic insights into the impact of key signaling pathways on lens populations at single-cell resolution. Fibroblast growth factor (FGF) signaling has been shown to play an important role in regulating epithelial to fiber differentiation in the lens^84^. Our own analysis in the present study has pointed to potential new aspects of FGF signaling in lens cell populations as indicated by, (1) progressive reduction of *FGFR3* in epithelial to fiber differentiation, and (2) transient upregulation of *FGFRL1* (FGF receptor like 1) across the epithelial to fiber trajectory. Therefore, we examined the impact of exogenous hyperstimulation of FGF on chicken lens development *in vivo*. The embryonic chicken eye has been used as an *in ovo* model for studying ocular development and retina regeneration^85–88^. We co-opted a retina regeneration paradigm in which the retina is surgically removed (retinectomy) and replaces with heparin beads that deliver FGF2 to the overlying lens (Supplementary Fig. S9). The lens was subjected to FGF2 treatment for 6 hours before sample collection for snRNA-seq. snRNA-seq data collected from retinectomy only (control) or retinectomy subjected to FGF2 hyperstimulation showed a dramatic change in cell populations (Fig. 9a). Using our previously-described developmental expression dataset as a reference (Fig. 2b), we then applied automated cell type annotation using a label transfer method to gain insights toward the cellular identity of these samples. Cells were categorized into epithelial, intermediate, and fiber states based on their transcriptional similarity to our developmental reference dataset. The control showed a similar spectrum of epithelial through fiber cell states, as previously observed (Fig. 9b; Fig. 3a). Interestingly, treatment with FGF2 beads results in a clear spatial separation of intermediate and fiber cell populations, which correspond to two novel clusters (Fig. 9c, Supplementary Fig. S10a). Further, quantification of cell assignments highlighted a clear increase in the proportion of FGF2-treated cells in intermediate and fiber cell states compared to control (Fig. 9d). Next, we examined expression of epithelial and fiber enriched genes in these populations. In the control, *TFAP2A* expectedly exhibits high expression in epithelial cells and reduced expression in fiber cells (Fig. 9e), while *ASL1* exhibits the opposite pattern (Fig. 9f). The relative expression of *TFAP2A* and *ASL1* reinforces the notion that FGF2 treatment results in the formation of a distinct intermediate and fiber population (Fig. 9e,f; Supplementary Fig. S10b).

**Fig. 9.**
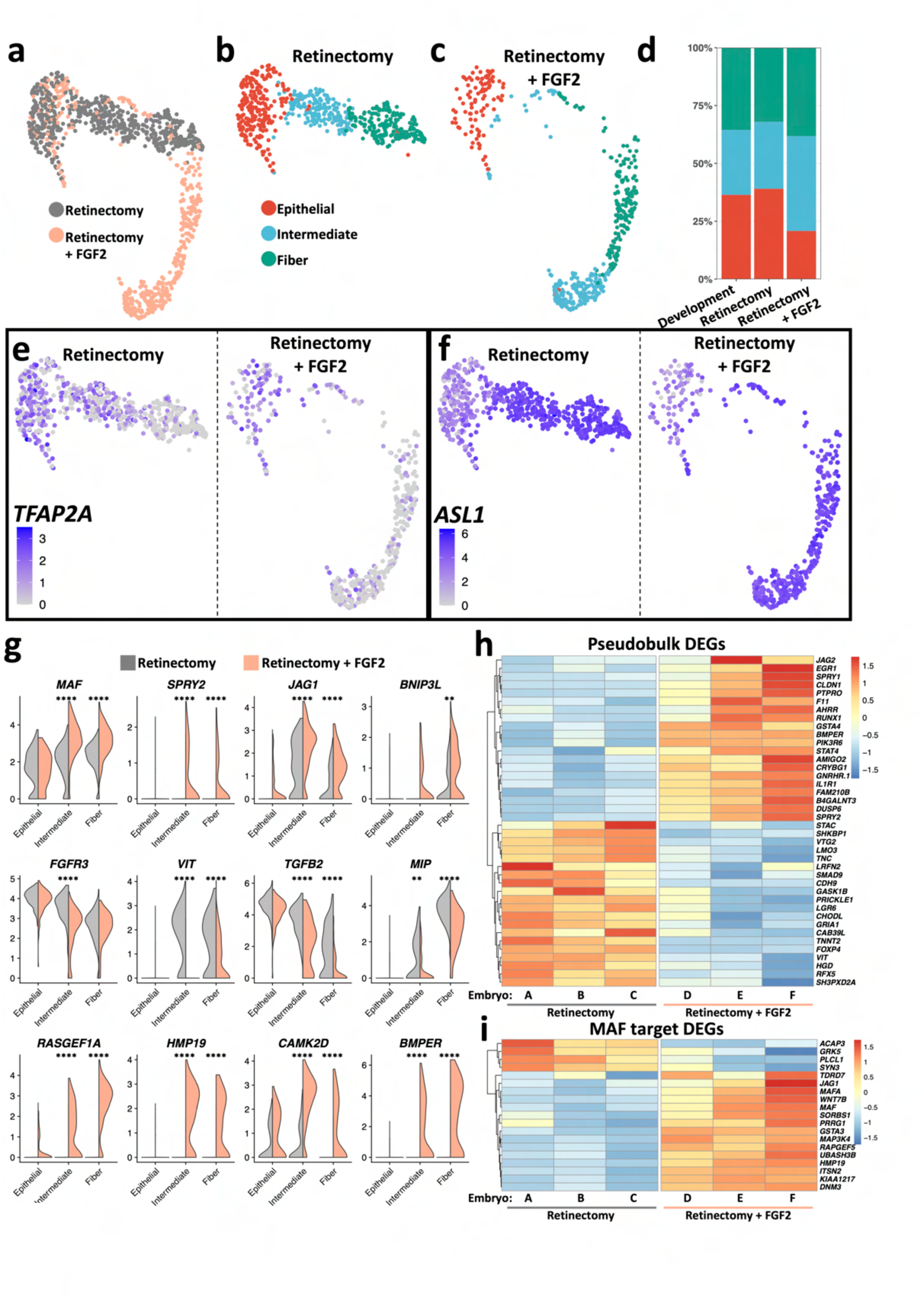
The FGF2-treated lens. (**a**): UMAP displays lens nuclei captured via snRNA-seq, performed using chicken eyes 6 hours after retinectomy ± FGF2-coated beads. (**b**): UMAP displays lens nuclei from the retinectomy-only (control) sample, colored by the predicted cell state when annotated against the development reference dataset. (**c**): Same as B, for FGF2-treated samples. (**d**): The proportion of nuclei assigned to each cell state for each condition and compared to the intact (developing) eye. (**e**): Feature plot displays log-normalized abundance of the epithelial-enriched gene *TFAP2A*; UMAP is split by condition. (**f**): Same as (e), but for the fiber-enriched gene *ASL1*. (**g**): Violin plots display the distribution of log-normalized transcript abundance for DEGs. ** denotes adjusted p-value < 0.005; **** denotes adjusted p < 0.00005; calculated via Wilcoxon Rank Sum test comparing retinectomy vs. FGF2-treated for each lens cell state. (**h**): Row-normalized heatmap displays select DEGs, calculated via pseudobulk analysis. Each column corresponds to a single individual (embryo). (**i**): Same as (h), displaying expression of predicted MAF regulatory targets. All genes in heatmap have adjusted p-value < 0.05.

After broadly defining FGF-responsive cell populations, we then focused on the altered expression of specific genes resulting from FGF treatment (Supplementary File S6). Several genes with a functional relation to FGF signaling (*e.g.*, *FGFR3*, *MAF*, *SPRY2*) were found to be responsive consistent with FGF hyperstimulation (Fig. 9g). Interestingly, *BNIP3L*, which is involved in fiber cell organelle degradation^89^, and in a separate study is down-regulated by FGF2 in H9C2 cells^90^, was found to be elevated in the present study. Paradoxically, while more cells adopted an overall intermediate identity in response to FGF2, the expression of *MIP* and *VIT*, which are genes associated with cataract or predicted to be important in the lens^11, 91^, was reduced relative to control (Fig. 9g). This could suggest that stimulation of fiber cell differentiation via exogenous FGF2 *in ovo* does not faithfully recapitulate the endogenous fiber differentiation program. This assay also identified genes that are upregulated upon FGF2 treatment (*e.g.*, *BMPER*, *RASGEF1A*; Fig. 9g). We then compared the expression of these genes to the endogenous expression patterns observed in the developing E4 lens (Supplementary Fig. S11). Consistent with expectations, we observed augmented expression of *MAF* uniquely in the FGF-stimulated condition, whereas *SPRY2* expression was extended into the fiber cell population following FGF2 stimulation. Similarly, relative to the developing lens, FGF2 stimulation resulted in suppression of *TGFB2* and *VIT* in intermediate or fiber cell populations. In addition, *BMPER*, *BNIP3L*, *CAMK2D*, *HMP19*, *JAG1*, and *RASGEF1A*, which are all expressed to some extent in the developing lens, were reduced in expression following retinectomy and in part recovered or increased in expression following FGF2 stimulation (Supplementary Fig. S11). Thus, it is possible that FGF2 is able to in part rescue the expression of certain lens genes that are not sustained in the absence of influences from the neural retina.

We next performed pseudobulk analysis, which summates total RNA across all lens cells in each condition, and we used this approach to uncover a cohort of genes responsive to FGF2 treatment (Fig. 9h). Aggregate differential expression analysis captured a clear treatment response (Supplementary Fig. S12a), identifying 471 genes significantly altered in lens cells (Supplementary File S6). In addition, this analysis captured numerous trends in gene expression that are consistent with a role of FGF2 in driving fiber cell differentiation, including elevated levels of *GATA3*, *CAPRIN2*, *PROX1*, *EPHA2*, *CDKN1B*, and *CDKN1C*, as well as reductions in *SIX3* and *PAX6*, although notably, not all of these genes reached statistical significance (adjusted p value > 0.05) via our pseudobulk approach (Supplementary Fig. S12b). Given the observed upregulation of *MAF* in response to FGF2, we next examined the overlap between FGF-responsive genes and those recognized as putative MAF targets (Fig. 6), which identified key genes linked to lens biology and defects, including cataract (*e.g.*, *JAG1*, *TDRD7*, etc.) (Fig. 9i). Together, these findings provide functional insights into the impact of FGF2-based activation of target genes in lens cells (Supplementary Fig. S13).

### Cataract-associated loci implicated in the present study

Our study captured potentially fundamental mechanisms of cell identity, differentiation, and homeostasis. We next sought to examine whether genes identified in these various approaches can serve to point toward the regulatory underpinnings of lens development and pathology. Therefore, we compared candidate genes identified in these various approaches with those identified in Cat-Map^92^ (an online databased of cataract-linked genes) and iSyTE^11, 20, 38^ (an approach to identify genes with lens-enriched expression that correlate with lens biology and pathology). The comparative analysis with Cat-Map identified 72 genes that are linked to cataract and serve as excellent markers of cell populations in the developing lens, spanning epithelial, intermediate, and fiber states (Fig. 10a; Supplementary File S7). We also identified 15 genes in Cat-Map that are identified as MAF targets, and a majority of which (80%) are expressed in fiber cells, which agrees with the previously-known, functional understanding of MAF’s role in the lens (Fig. 10b). Analysis of PRC2 targets that are found in Cat-Map suggested that few PRC2-bound genes are associated with lens cell populations; the majority of these genes are not significantly regulated or not appreciably expressed in lens cells at the examined developmental stage (Fig. 10c). This is consistent with the regulatory logic of PRC2 as a repressor and may potentially explain the contribution of these cataract-linked genes to lens pathology, which may result from spurious expression secondary to defective transcriptional silencing. Similarly, numerous FGF2-responsive genes were identified to have an overlap with Cat-Map (Fig. 10d). Interestingly, a number of FGF2 targets were not robustly expressed or associated with specific cell populations in our development-only dataset; this could point toward dysfunctional FGF-signaling as a mechanism for aberrant gene activation and subsequent cataractogenesis. Analysis of the top 528 genes with lens-enriched expression in the iSyTE database followed a similar trend (Supplementary Fig. S14; Supplementary File S7). Together, these analyses identify new genes associated with lens development, homeostasis, and pathology.

**Fig. 10.**
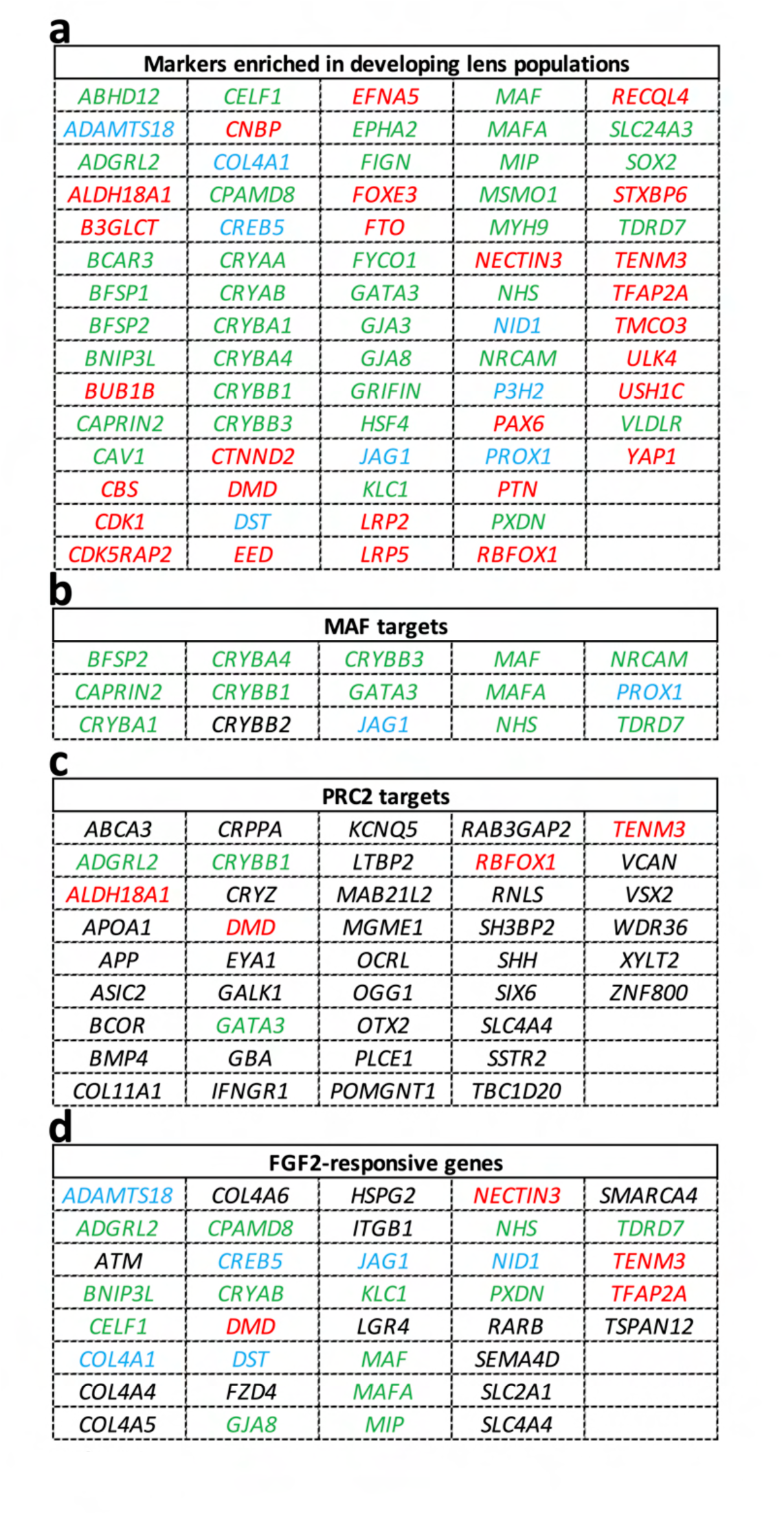
Cataract-associated genes. Tables summarize the genes identified in the current study that are documented in the CAT-MAP database, a repository for cataract-associated genes https://cat-map.wustl.edu/. Lists encompass significant marker genes drawn from the developing chicken snRNA-seq dataset (**a**), genes with regulatory correlation to nearby predicted MAF binding sites (**b**), genes bound by PRC2 in the developing chick lens (**c**), and genes with significant regulation in response to FGF2 hyperstimulation (**d**). Genes are colored according to their enrichment patterns during lens development: Red = Epithelial-enriched; Blue = Intermediate-enriched; Green = Fiber-enriched; Black = Not significantly enriched in a cluster during development, or not expressed.

## Discussion

While recent studies have reported single-cell RNA-seq analysis of the lens in zebrafish^52^, mouse^53^, and human^93–95^, there remains a need to apply single-cell multiomic outlooks to comprehensively examine lens development. The present work encompasses a comprehensive multiomic analysis of the embryonic chicken lens at single-cell resolution, which portrays a novel, holistic perspective of epithelial to fiber cell differentiation and the associated gene regulatory mechanisms. Briefly, we performed snRNA-seq, snATAC-seq, and CUT&RUN-seq, as well as a functional assay (FGF2 hyper-stimulation) during a critical, early window of chicken lens development. To contextualize these findings and gain new biological insights, we carried out an integrative analysis in the context of other information relevant to lens biology, such as motif analysis for key lens TFs (*e.g.*, FOXE3, MAF), and related these findings to cataract-linked genes in Cat-Map^92^ and high-priority lens genes in the iSyTE database^11, 20, 38^. Specifically, this report: (1) defined various single-cell regulatory states in the lens, in turn providing a detailed spatiotemporal roadmap of epithelial to fiber cell differentiation, (2) uncovered key regulatory events underpinning lens development, including important pathways and TFs, as well as intrinsic and extrinsic cues governing this process, (3) captured the dynamic reconfiguration of lens chromatin, including the *cis*- and *trans*-regulatory events that control fiber cell differentiation, (4) identified PRC2-directed epigenetic surveillance as a novel mechanism for cell fate control in the lens, (5) assessed the functional impact of FGF hyper-stimulation on lens cell identity by profiling the emergence of novel lens cell states and the activation of our predicted MAF downstream network, and finally, (6) provided a developmental and/or mechanistic frame of reference for cataract-associated genes, thereby prioritizing novel effectors of lens development as they relate to pathology.

Cumulatively, this study represents a model for applying multiomic approaches and integrative analysis to tissue development. We demonstrated how these efforts led to the identification of novel, conserved mechanisms of lens development, and we further set the stage for capturing fundamental processes of cellular differentiation and pathology. A recent review proposed the implementation of various bulk assays, such as RNA-seq, ATAC-seq, and CUT&RUN-seq, that would stand to advance our understanding of mechanisms central to lens biology^51^. Our study builds on these approaches by capturing multiomic readouts from the same sample (combined snRNA/snATAC), and further represents a seminal first step toward applying these principles to the lens at single-cell resolution. Even across vertebrate development, our study is among the few to apply CUT&RUN-seq to effectively profile complex features of DNA binding proteins and transcription regulators *in vivo*. We contend that these complimentary, multimodal approaches provide unified insights toward specific mechanisms of development, such as the complex regulation of *GATA3* activation in the lens epithelium or the nature of PRC2-chromatin regulatory dynamics.

In addition to identifying PRC2 as novel regulatory protein complex in the lens, we also demonstrate how PRC2 is involved in controlling the dynamic expression of genes encoding key lens TFs, such as *GATA3*, which itself has an important role in fiber differentiation. While PRC2 has been shown to be involved in the formation and maintenance of heterochromatin^60–62^, we demonstrate presently, for the first time, the participation of this key regulatory complex in lens development *in vivo*. Based on the striking distribution of PRC2 protein components at specific loci across the lens developmental genome, it can be postulated that this complex may be involved in the active suppression of alternate cell fates in the lens, including non-lens ocular lineages. For example, retinal genes such as *ASCL1*, *RAX*, *SIX6*, *VSX2*, among others, exhibit a high degree of PRC2 binding and H3K27me3 localization signal, pointing toward their active suppression by PRC2 in lens cells. Indeed, other studies have suggested that lens cells may have a predisposition for acquiring aberrant retinal-like transcript profiles, and documented perturbed H3K27 trimethylation levels linked to this phenotype^96^. Our data now show that a PRC2-mediated regulatory mechanism may be involved in suppression of retinal cell fates in the lens. In an analogous context, chromatin proteomics was used to associate PRC2 with lineage control in cultured human embryonic stem cells^97^. Our study utilizes CUT&RUN to resolve specific PRC2-binding events to demonstrate that this PRC2-based mechanism is also co-opted later in development, in a tissue-specific manner to control cell fate. Future studies can be designed to assess whether lens phenotypic plasticity can be augmented through the perturbation of PRC2 activity. Interestingly, we observe the loss of core PRC2 subunits and H3K27me3 as fiber cells differentiate, concurrent with progressive increases in *JARID2* abundance. While the present study does not resolve the functional significance of these PRC2 dynamics, it can be speculated that loss of PRC2 may reflect a hallmark of terminal differentiation. Whether these changes in PRC2 set the stage for later facets of fiber cell differentiation, such as organelle degradation, remains to be investigated.

Our study further attempts to leverage high-content multimodal information to reconstruct complex features of lens gene regulation, exemplified by the prediction of regulatory outputs for the human cataract-linked TF MAF. With this approach, we not only identify previously known MAF targets, but further expand the connectivity of this key TF by predicting new regulatory targets of MAF in the lens. It should be acknowledged that the *in-silico* deduction of gene regulatory networks is an emerging field experiencing active software development^98–101^, and accordingly, some regulatory relationships may need to be independently validated. In this regard, *in-silico* regulatory predictions can miss scenarios where chromatin accessibility is not dramatically affected, or where TFs bind DNA indirectly. It can further be challenging to disambiguate between TFs that bind to closely related motifs, such as those within the same protein family. The advent of systems-level multiomic approaches, such as combined snRNA/snATAC-seq, stand to better refine these predictions by establishing complementary layers of evidence that can reinforce the observed regulatory relationships. As the field of *in-silico* prediction of gene regulatory networks expands, this predictive approach can be applied to other factors, *e.g.*, *FOXE3*, *HSF4*, *MAB21L1*, *PITX3*, *TFAP2A*, etc., for which loss-of-function global gene expression datasets have been generated and thus can be used to validate predictions^102–108^.

Finally, FGF signaling is known to be a driver of lens fiber differentiation^84, 109^, and our present study reinforces its role from the perspective of single-cell dynamics. In this regard, following FGF2 stimulation, we observe an overall reduction in the proportion of epithelial cells, as well as altered intermediate and fiber cell states relative to our retinectomy-only control. Furthermore, we document the activation of *MAF* in an FGF-dependent manner *in vivo*, reinforcing MAF as a target of FGF^110^. Our predictive model places MAF as a key player in driving fiber cell differentiation, as we demonstrate that FGF-hyperstimulation results in the accelerated activation of MAF and the upregulation of several predicted MAF targets. Given that several genes are reduced in expression following retinectomy and recovered in the presence of FGF2, it could be tempting to speculate that the neural retina serves as an essential source of certain growth factors, which can be in part rescued by exogenously supplied FGF2. In summary, our study leverages numerous nascent omics approaches to intertwine a global perspective of gene regulation with granular insights, serving to reinforce known regulatory mechanisms in the developing lens and extend our understanding into previously unappreciated facets of cellular differentiation and organogenesis.

## Methods

### Chicken embryos and nuclei isolation

Fertile Specific Pathogen Free eggs (Charles River Laboratories, 10100329) were incubated in a rotating, humidified incubator at 37° C. Whole chicken eyes were enucleated at E4 or E5 in PBS containing 0.2 U / µl RNasin Plus RNase Inhibitor (Promega, N2615). Eyes from 3 embryos were pooled for each condition. Nuclei were immediately extracted and fixed for snRNA-seq. For combined snRNA-seq and scATAC-seq, samples were first cryopreserved via snap freezing in an isopentane bath on liquid nitrogen and briefly stored at -80°C prior to nuclei extraction. In brief, nuclei extraction was carried out via an optimized variation of the demonstrated protocols described in 10X Genomics Protocol CG000124 Rev E and 10X Genomics Protocol CG000366 Rev D. For snRNA-seq, cell lysis was performed by triturating sample and incubating in a cold buffer comprised of 10 mM Tris-HCl, 10 mM NaCl, 3 mM MgCl_2_, 0.01% Surfact-Amps NP-40 (Thermo Fisher Scientific, 28324), and 0.2 U / µl RNase Inhibitor. Nuclei were washed 3 times by resuspending in cold PBS containing 2% fraction V bovine serum albumin and 0.2 U / µl RNase inhibitor (Promega, N2615). For multiome assay, lysis was performed via trituration and 30 second incubation in a buffer containing 10 mM Tris-HCl, 10 mM NaCl, 3 mM MgCl_2_, 0.005% Tween-20, 0.005% Surfact-Amps NP-40, 0.0005% digitonin (Thermo Fisher Scientific, BN2006), 1% fraction V bovine serum albumin, 1 mM DTT, and 1 U / µl Sigma Protector RNase inhibitor (MilliporeSigma, 3335402001). Nuclei were washed twice by resuspending in cold buffer containing 10 mM Tris-HCl, 10 mM NaCl, 3 mM MgCl_2_, 1% fraction V bovine serum albumin, 0.1% Tween-20, 1 mM DTT, and 1 U / µl Sigma Protector RNase inhibitor. Nuclei were passed twice through 20 µm cell strainers (pluriSelect, 43-10020-40) during isolation to filter debris and cell aggregates.

### Single nucleus multiome profiling library preparation and data generation

Nuclei were subjected to the Chromium Next GEM Single Cell Multiome ATAC + Gene Expression library preparation workflow (10X Genomics, 1000285 & 1000230) following 10X Genomics protocol CG000338 Rev F, loading approximately 16,000 nuclei per reaction. The 10X Genomics Chromium Controller was used for GEM generation. ATAC libraries were amplified with 7 total PCR cycles, and RNA libraries were amplified with 15 PCR cycles. Indexes were incorporated from Single Index Kit N, Set A (10X Genomics, PN-1000212) and Dual Index Plate TT Set A (10X Genomics, PN-1000215). Libraries were sequenced at the Novogene Sequencing Core (Sacramento, CA) on the Illumina NovaSeq 6000. ATAC libraries were sequenced to generate 49,000-54,000 reads per nucleus and RNA libraries were sequenced to generate 29,000-56,000 reads per nucleus.

### Split-pool barcoding library preparation and data generation

snRNA-seq libraries were generated via the split-pool barcoding method^111^ using the Parse Biosciences Single Cell Whole Transcriptome Kit Chemistry Version 1 (Parse Biosciences, SB2001). Nuclei were fixed according to the Nuclei Fixation Kit Protocol (Parse Biosciences, SB1003) and stored at -80°C. During library preparation, cDNA was amplified with 13 total PCR cycles. Final libraries were assessed with the Agilent Bioanalyzer and Qubit 4. Libraries were sequenced across two lanes of HiSeq X Ten and a lane of NovaSeq 6000 at the Novogene Sequencing Core (Sacramento, CA) to generate an average of 48,101 reads per nucleus across 58,906 total recovered nuclei.

### Single-nuclei sequence alignment, gene annotation, and data pre-processing

Downstream analysis of single-nuclei data was performed using command line interface, R version 4.2.2, and RStudio Version 2023.06.0+421. Reads were aligned to chicken genome GRCg7b and analyzed using the chicken annotation from Ensembl Release 109^112^. Throughout the analysis, processing of the chicken annotation was facilitated using the R package ensembldb (v2.22.0)^113^. For comparative studies, orthology was determined using Ensembl Biomart. Ensembl gene ENSGALG00010026936 was designated as *FOXE3* throughout the manuscript. Reference genome indexes were built for the alignment of single-cell data using the Parse Biosciences alignment suite (v1.0.3) and 10X Genomics Cell Ranger ARC (v2.0.2). Filtered gene matrices were read into the R environment using Seurat (v4.3.0)^114^. Nuclei with 200 > genes, < 6000 genes, < 10% mitochondrial reads, and < 15000 RNA molecules were retained in the Parse Biosciences dataset. Nuclei with 200 > genes, < 5000 genes, < 15% mitochondrial reads, and < 15000 RNA molecules were retained in the 10X Genomics dataset. Ambient RNA was subtracted from raw reads using the SoupX package (v1.6.2)^115^, with the estimated contamination fraction set to the value calculated via Souporcell, as described below^116^.

### In-silico genotyping and doublet removal

*In-silico* genotyping was employed to achieve biological replication by assigning chicken nuclei, derived from 3 pooled embryos per condition, back to an embryo of origin. Genotyping was facilitated by the Souporcell software (v2.0)^116^, using either the provided singularity or on a step-by-step basis. The detailed workflow for Souporcell was followed as written (https://github.com/wheaton5/souporcell). This workflow relies on additional software, including Samtools^117^, Minimap2^118^, and Freebayes^119^. Single nucleotide variants were cataloged using chicken genome GRCg6a for Parse Biosciences data and genome GRCg7b for 10X Genomics data. Prior to genotyping, cell barcodes in bam alignment files were given unique suffixes for each library and merged. Barcode assignments were read into the R environment and integrated into Seurat objects. Nuclei labeled as doublets, as well as a small number of nuclei with invalid genotype assignments (nuclei assigned to a condition incompatible with their sample of origin), were discarded from the analysis.

### snRNA clustering, marker identification, differential expression testing, and pathway enrichment analysis

Filtered nuclei datasets were processed with Seurat^114^, relying on sctransform for normalization and variance stabilization^120^ prior to clustering nuclei and UMAP visualization. Cell cycle scoring was performed using the Seurat *CellCycleScoring* function, and the difference between S phase score and G2M Score was saved and used as a regression variable during normalization. In addition, the percentage of counts assigned to the mitochondrial and W chromosomes were calculated and regressed during normalization. Analysis of each dataset resulted in clear nuclei populations expressing *ASL1* that were subset and re-processed for downstream analysis. These lens populations were integrated by applying the *PrepSCTIntegration* function, followed by anchor-based integration. The integrated lens dataset was subjected to PCA, UMAP, and nearest neighbors analysis with 25 dimensions of reduction and clusters, determined using Seurat *FindClusters* and Louvain algorithm, with the resolution parameter set to 0.2. Raw counts were log normalized and were used for downstream differential expression testing, marker identification, and visualization, unless otherwise specified. Differential expression testing between clusters of nuclei was performed using the Wilcoxon Rank Sum test (Supplementary File S1). Pathway enrichment analysis was performed using gprofiler2^121^, using either the R package (v0.2.1) or online interface (Supplementary File S2). Gene lists were converted to human orthologs and run as multi-query assays for pathway enrichment analysis.

### Pseudotime and trajectory analysis

Pseudotime trajectories were inferred using the Monocle 3 software (v1.3.1)^122–124^. Seurat objects were converted to Monocle cell data sets using the SeuratWrappers package (v0.3.1) function *as.cell_data_set*. Pseudotime trajectories were preprocessed using the Monocle workflow, including normalization, clustering, graph learning, and cell ordering^125, 126^. Twenty-five dimensions were applied during processing and the *minimal branch length* parameter was adjusted to 5 during graph learning. Moran’s I test was applied for differential expression testing across the trajectory using the principal graph and an adjusted p-value cut-off of 0.05 was applied (Supplementary File S3). Expression across the trajectory was visualized using the *plot genes in pseudotime* function with parameters *min expr = 1* and *trend formula = “∼ splines::ns(pseudotime, df=3)”*. Expression heatmaps were generated using the pheatmap package (v1.0.12) with columns ordered based on the pseudotime ranking assigned from the pseudotime analysis. For heatmaps, the rolling mean of log normalized counts was calculated using the *rollmean* function of the R package zoo (v1.8-11)^127^, using the neighboring 5% of cells as a rolling window.

### ATAC-seq data processing, peak calling, and differentially accessible region testing

ATAC data output from Cell Ranger ARC was read into R as a merged hdf5 file using Seurat. Fragments were added to the existing RNA-only file and Multiome nuclei were subset for further analysis. Nucleosome signal and TSS signal was calculated using Signac^128^ and nuclei were retained which contained less than 40000 ATAC counts, greater than 1000 ATAC counts, nucleosomal signal less than 2, and TSS enrichment greater than 1. Peak regions were defined using HMMRATAC^129^ (v1.2.10), using the Cell Ranger ARC bam alignment file filtered to contain only lens nuclei. Peaks with a score < 10 were removed and mitochondrial peaks discarded. ATAC counts were normalized with the term frequency inverse document frequency method^130^ using the *RunTFIDF* function. Partial singular value decomposition was performed with the Signac (v1.9.0) *RunSVD* function and UMAP was generated using the latent semantic indexing reduction. The combined modality UMAP was generated using the Seurat function *FindMultiModalNeighbors* to generate a weighted nearest neighbors graph derived from the principal component analysis reduction generated for RNA and the latent semantic indexing reduction generated for ATAC. Differential accessibility testing between clusters was performed using the logistic regression and likelihood ratio “*LR*” test in Seurat with the *min.pct* parameter set to 0.05. Coverage plots were generated using the Signac *CoveragePlot* function to display normalized Tn5 insertion frequency across loci of interest.

### Linkage of accessible regulatory elements to genes, motif profiling, and chromVAR analysis

The correlation of peak accessibility with nearby gene expression was carried out using the Signac *LinkPeaks* function^131^, which takes into account GC content and peak length. Normalized RNA and ATAC values were used as input, Peak-TSS relationships were confined to 1000000 base pairs, and a p-value cut-off of 0.05 was applied. Peaks were annotated with vertebrate motifs drawn from the 2022 JASPAR CORE database^72^. Motifs were added to the Seurat object using the Seurat *AddMotifs* function and visualized. chromVAR analysis^132^ was carried out using the Signac function *RunChromVAR* and adjusted p values were determined by applying a Wilcoxon Rank Sum test to motif activity scores. Motif logo displays were made using the Signac *MotifPlot* function. For the prediction of MAF binding events, the prediction of transcription factor binding events was drawn from the annotated Seurat objects. Predicted peak-TSS linkages were retained which both 1) contain a MAF motif in the peak region of and 2) exhibit a positive correlation coefficient between peak accessibility and gene expression, which was reasoned based on the accepted activity of MAF as a transcriptional activator^133^. The rolling mean of normalized RNA abundance and accessibility were then calculated across the pseudotime trajectory, as described above. Peaks with a normalized accessibility that exhibited a positive correlation with *MAF* RNA abundance (correlation coefficient > 0.6) were retained as candidate MAF-binding loci.

### Hybridization chain reaction fluorescent in-situ hybridization

Fertile chicken eggs were obtained from either Michigan State University or Charles River Laboratory (10100329). Embryos were rinsed in PBS, fixed overnight in 4% PFA at 4°C, washed in PBS, and stored in 100% MeOH at -20°C. HCR-FISH was carried out using custom probes and reagents designed by Molecular Instruments (Los Angeles, CA) and a modified version of the Molecular Instruments protocol *MI-Protocol-RNAFISH-Mouse Rev. 8 2022-07-22* (https://www.molecularinstruments.com/hcr-rnafish-protocols) was used. In brief, samples were rehydrated in graded MeOH / PBST washes and permeabilized with 10 µg / ml proteinase K for 20 minutes prior to 20 minutes of post-fixation with room-temperature 4% PFA. Sets of 20 custom probes each against ENSGALT00010019926.1 (*EZH2*) and ENSGALT00010023105.1 (*JARID2*) were introduced at 2 pmol and incubated overnight at 37°C. Hairpin incubation was performed overnight at room temperature using 60 nM of each hairpin. Samples were briefly equilibrated in 30% sucrose at 4°C before snap freezing in optimal cutting temperature (OCT) medium on dry ice. Samples were sectioned on a cryotome, stained with 1 µg/ml DAPI (MilliporeSigma, MBD0015), washed, and a coverslip applied with Fluoromount prior to imaging. Imaging was performed using the Zeiss LSM 710 Laser Scanning Confocal System using the Zen Black image software, and post-processing of images was done using the Fiji ImageJ platform^134^. Images shown are representative of at least 3 embryos.

### Immunohistochemistry

*FVB/N* mouse embryos were collected at embryonic day 12.5 or 14, as specified, and fixed overnight in 10% formalin at 4°C prior to paraffin embedding and sectioning. Immunostaining was performed as described previously^135^. In brief, sections were deparaffinized and antigen retrieval was performed at 100°C for 15 minutes with 0.01M sodium citrate, pH 6.0. Sections were permeabilized in 1% saponin for 5 minutes. Blocking was performed with 10% goat serum. Samples were incubated with primary antibodies at the specified dilutions for 4 hours at room temperature; secondary antibodies were incubated for 2 hours at room temperature. Images shown are representative of at least 2 embryos. Antibodies and dilutions employed in this study are summarized (Table 1). Imaging was performed using the Zeiss LSM 710 Laser Scanning Confocal System using the Zen Black image software, and post-processing of images was done using the Fiji ImageJ platform^134^.

**Table 1.**
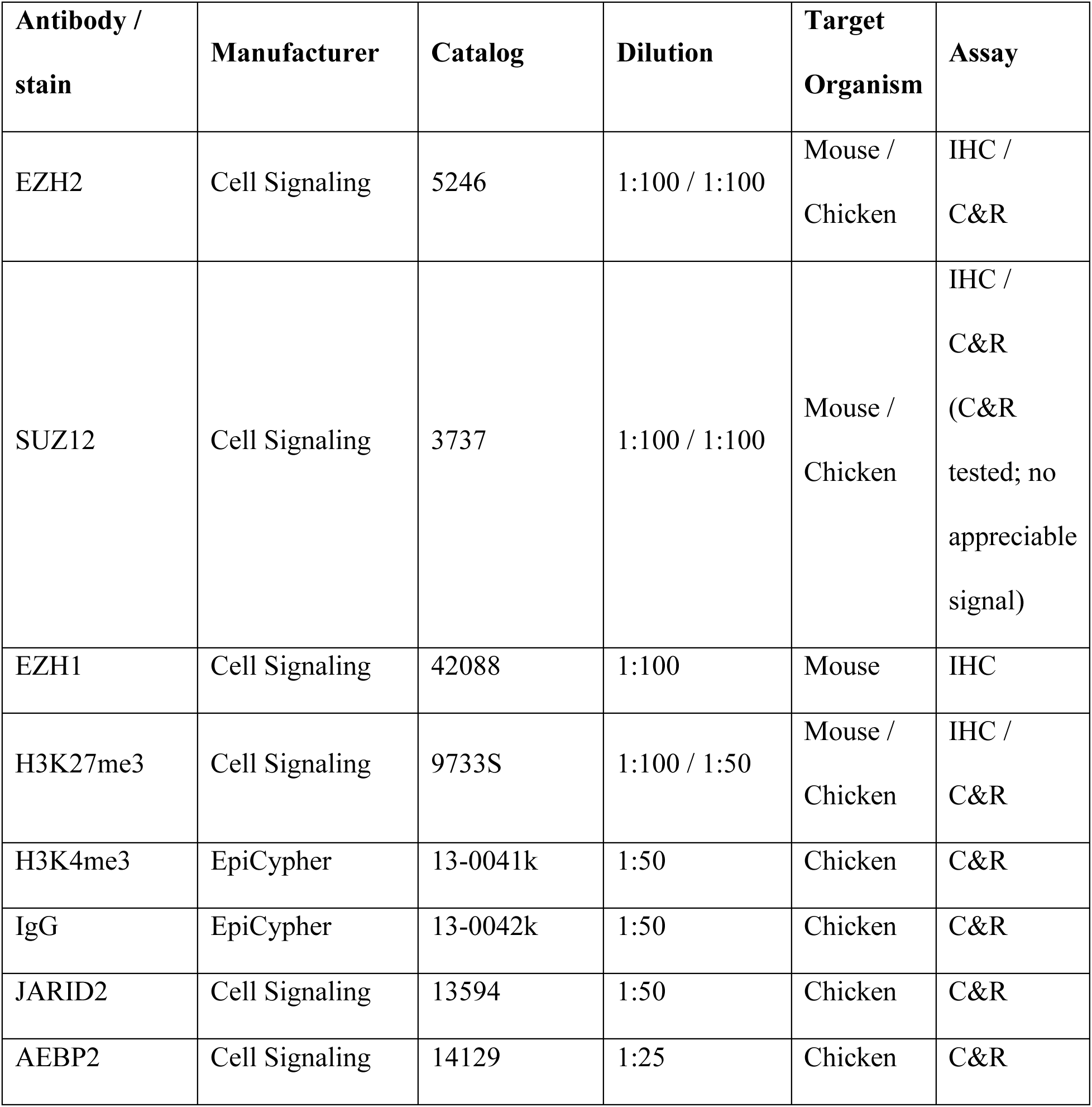
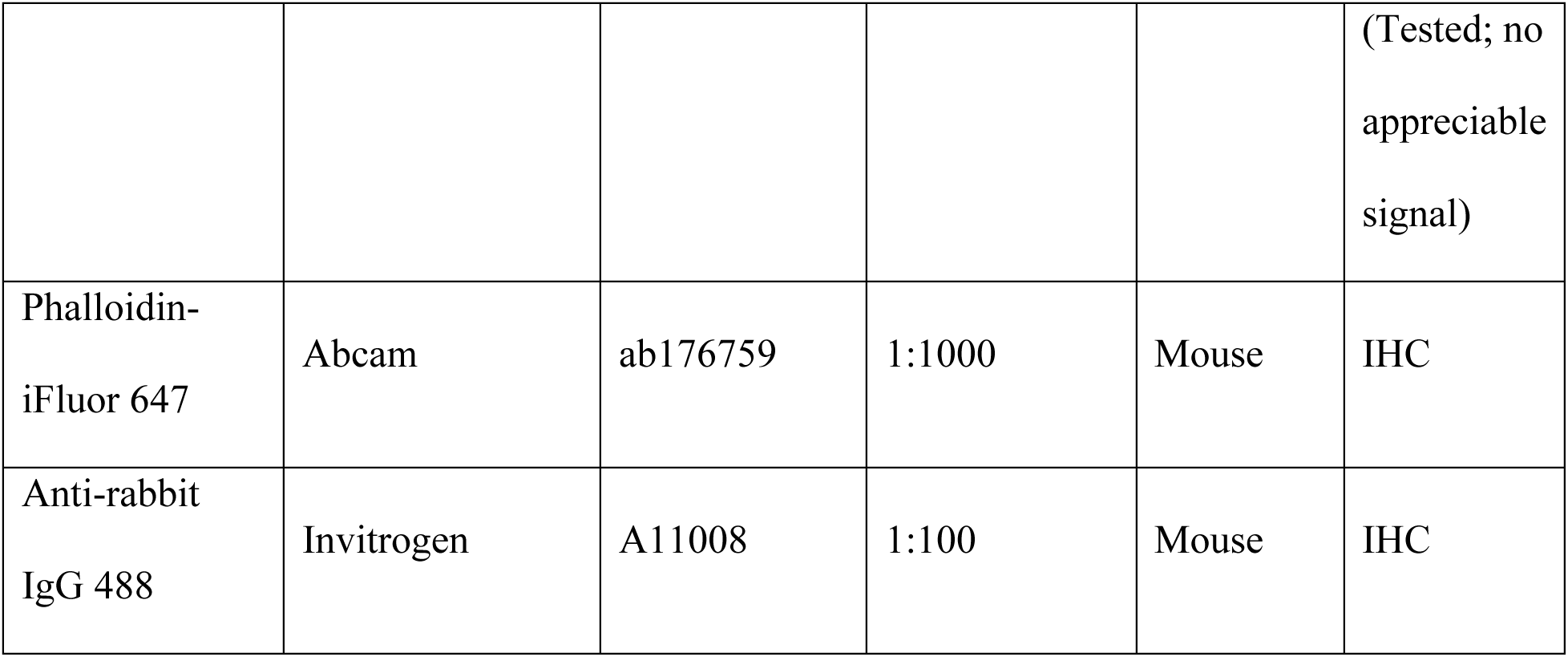
Summarizes antibodies and stains used in the study. IHC = immunohistochemistry; C&R = CUT&RUN-seq.

### CUT&RUN-sequencing and analysis

Cleavage Under Targets and Release Using Nuclease sequencing (CUT&RUN-seq)^83, 136, 137^ was performed using the CUTANA ChIC / CUT&RUN Kit Version 3 (EpiCypher, 14-1048) and CUTANA CUT&RUN Library Prep Kit (EpiCypher, 14-1002). The protocol was carried out according to manufacturer protocol, with slight modification. Chicken embryos were obtained from Michigan State University and lenses were dissected from embryonic day 4.5 chicken embryos. Dissected lenses were collected on ice and pooled into 3 biological replicates consisting of approximately 44 lenses each. Nuclei were extracted from samples using the protocol described above for single-nuclei applications. Buffers were modified to contain 1X proteinase inhibitor (MilliporeSigma, 11873580001), 1.5 mM spermidine, no RNase inhibitors, no DTT, and lysis buffer was modified to contain 0.001% Surfact-Amps NP-40. Nuclei across all samples were diluted to 1000 nuclei / µl and loaded at 100000 nuclei per reaction. All antibody reactions were performed in triplicate across independent biological replicates, and the negative control IgG antibody reaction was performed in duplicate. Antibodies employed in this study are summarized (Table 1; Supplementary File S8). Final libraries were assessed on the Agilent Bioanalyzer and Qubit 4 before pooling and sequencing at the Novogene Sequencing Core (Sacramento, CA) on a lane of Illumina HiSeq X Ten. Raw reads were quality and adapter trimmed using Trim Galore! (v0.6.4_dev)^138^ with parameters *–phred33 –paired –length 36 -e 0.1 -q 5 –stringency 1*. Trimmed reads were aligned against chicken genome GRCg7b with Bowtie 2 (v2.4.1) using parameters *-X 2000 --local --no-mixed --no-discordant --no-dovetail*. Peak calling and normalization of CUT&RUN-seq data was performed similar to as described^139^, with the following modifications. Peaks were called using MACS2 (v2.2.7.1)^140^ and parameters *-f BAMPE -g 1070887886 --keep-dup all -q 0.05*, providing IgG alignment files as background. A minimum integer score of 400 was set for defining EZH2 peaks and 125 for defining JARID2 peaks. Each sample was normalized prior to visualization using the deepTools^141^ *bamCoverage* (v3.5.1) function with parameters *--binSize 1 --normalizeUsing RPGC --effectiveGenomeSize 1070887886*. RPGC normalization (reads per genomic content) corresponds to 1x normalization. Normalized replicates were merged into a single file for visualization. CUT&RUN heatmaps were produced using the deepTools *computeMatrix* (v3.5.1) and *plotHeatmap* (v3.5.1) functions using merged bigwig files. EZH2 and JARID2 co-localization was determined with the GenomicRanges R package (v1.50.2)^142^. The annotation of peak regions to genomic elements was performed with the ChIPseeker (v1.34.1) R package^143^.

### Chicken surgeries and FGF2 delivery

Fertile specific pathogen-free chicken eggs were obtained (Charles River Laboratories, 10100329). Retinectomies and FGF2 delivery were performed at embryonic day 4 as previously described^87, 144^. In brief, an incision was made around the anterior eye chamber with surgical scissors and the neural retina dislodged with a wire probe. For FGF2-delivery, approximately 20 acrylic beads with immobilized surface heparin (MilliporeSigma, H5263) were incubated overnight in 4 µl of 250 ng/µl of bovine basic FGF2 (R&D Biosystems, 133-FB-025). At the time of surgery, FGF2 beads were deposited into the posterior chamber of the eye cup and the eye shut. Embryos were incubated for 6 hours before enucleating the eyes for use with the Parse Biosciences snRNA-seq workflow, as described above. One eye from 3 chickens was pooled per condition.

### Analysis of FGF2-treated samples

Pre-processing of snRNA-seq data for FGF2-treated samples was performed as described above for development samples. Using this approach, a multiplet lens population was identified that exhibited abnormally high total RNA counts and total genes detected, as well as co-expression of numerous retinal and mesenchyme-specific genes (Supplementary Fig. 15), and this population was discarded from downstream analysis. FGF2-treated lens cells were assigned cluster labels using the Seurat *TransferData* function, using the Parse Biosciences-derived nuclei from the development dataset as a reference. Pseudobulk analysis was performed by summation of the ambient RNA-adjusted raw counts associated with each embryo, and sums were rounded to the nearest whole integer. Genes with < 10 total counts, as well as genes derived from the W chromosome, were discarded from the count matrix prior to analysis. Normalization and differential expression testing were performed using DESeq2 (v1.38.3)^145^.

## Supplementary files

*Supplementary File S1*.

Displays gene markers identified for lens cell populations in snRNA-seq analysis.

*Supplementary File S2*.

Summarizes pathway enrichment results employed throughout the study.

*Supplementary File S3*.

Contains statistics for detecting genes which vary across the pseudotime trajectory.

*Supplementary File S4*.

Video displays H3K27me3 localization in the E12 mouse lens.

*Supplementary File S5*.

File summarizes the localization of EZH2 and JARID2 peaks detected via CUT&RUN-seq.

*Supplementary File S6*.

File contains all differential expression testing results from FGF2 experiment.

*Supplementary File S7*.

File summarizes overlapping genes between the present study and CATMAP and iSyTE databases.

*Supplementary File S8*.

Contains details related to CUT&RUN library preparation.

## Acknowledgements

JAT was supported by National Institute of Neurological Disorders and Stroke [F99 NS129167]. KDRT was supported by National Eye Institute [R01 EY026816], the Rapid Grant Program at Miami University, and the John W. Steube Endowed Professorship. S.A.L. was supported by National Institutes of Health / National Eye Institute [R01 EY021505 and R01 EY029770]. M.L.R. was supported by National Institutes of Health / National Eye Institute [R21 EY033471 and R21 EY031092]. SR was supported by the “Choose Development!” Program from the Society for Developmental Biology and the Miami University Chapter of the Louis Stokes Alliance for Minority Participation Research Grant.

The authors thank Carlos M Charris Dominguez for assistance with nuclei isolation and Anil Upreti for valuable discussions throughout the execution of this study. We acknowledge and thank the staff (Dr. Andor Kiss & Ms. Xiaoyun Deng) of the Center for Bioinformatics & Functional Genomics (CBFG) at Miami University for instrumentation and computational support. The authors further acknowledge the use of Miami University’s Redhawk HPC cluster and thank Dr. Jens Mueller for computational support. The Miami University Center for Advanced Microscopy and Imaging (CAMI) supported imaging throughout this manuscript, including support from Dr. Zach Oestreicher and Mr. Matt Duley. Dr. Andy Fischer and Dr. Heithem El-Hodiri of The Ohio State University generously assisted with multiome library preparation. Jackson Theile assisted with the design of several schematics used throughout the manuscript.

## Author Contributions

JAT performed library preparation and sequence data generation. JAT and SB performed bioinformatic analysis. JAT, SMR, EG, and JMR contributed to sample collection and imaging. JAT and SAL were primarily responsible for writing, with all authors editing the final manuscript. JAT, MLR, SAL, and KDRT were primarily responsible for the study conception, experimental design, and data interpretation.

---

**Supp. Fig. 1.**
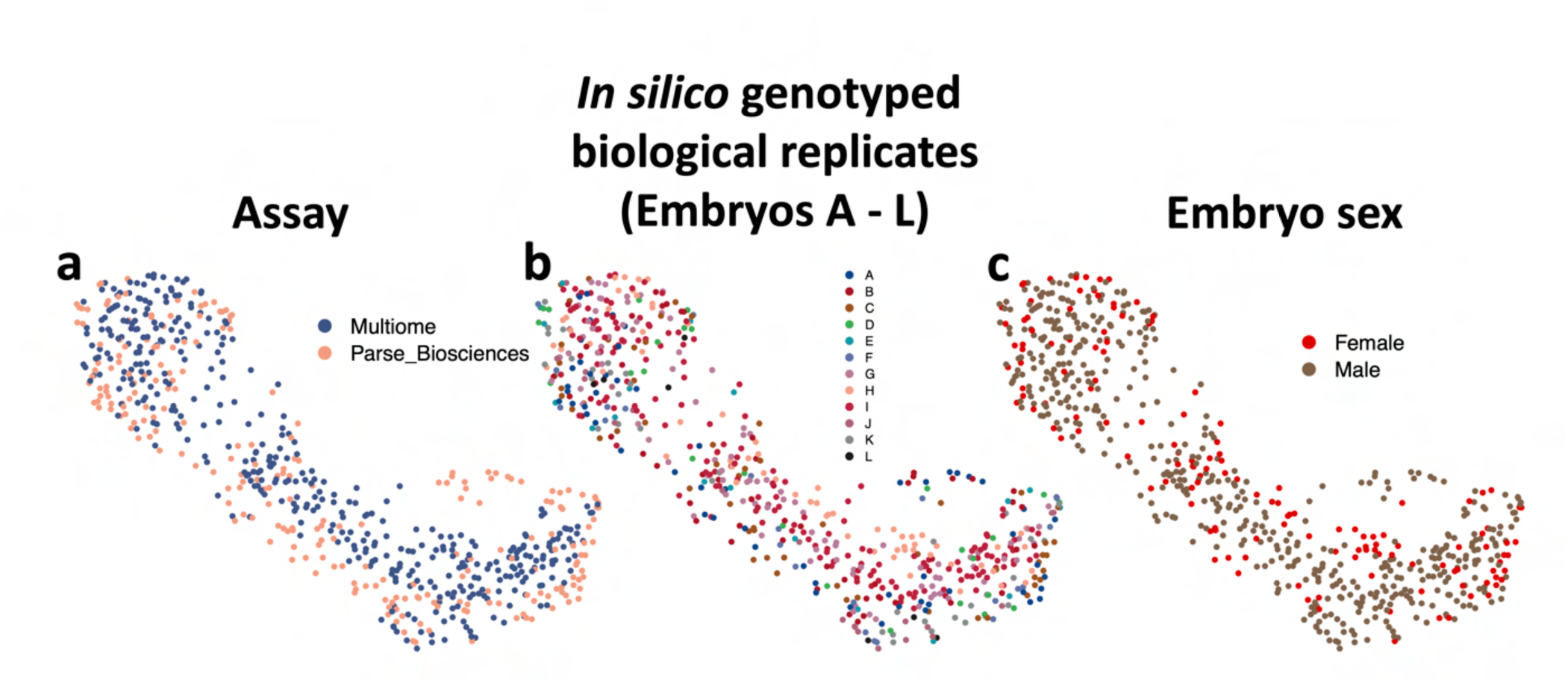
**(a):** UMAP represents captured lens nuclei transcriptomes colored by library preparation method: either lOX Genomics Multiome assay or Parse Biosciences whole transcriptome. **(b):** UMAP is color coded by biological replicate. 3 biological replicates were pooled for each run across 2 time points (E4 / ES) and 2 library preparation methods (12 embryos total). Embryo of origin was deduced by *in silico* genotyping and embryos were assigned an arbitrary letter A-L. **(c):** The sex of each embryo was deduced by assessing the RNA abundance levels of genes on the W chromosome and plotted on a UMAP.

**Supp. Fig. 2.**
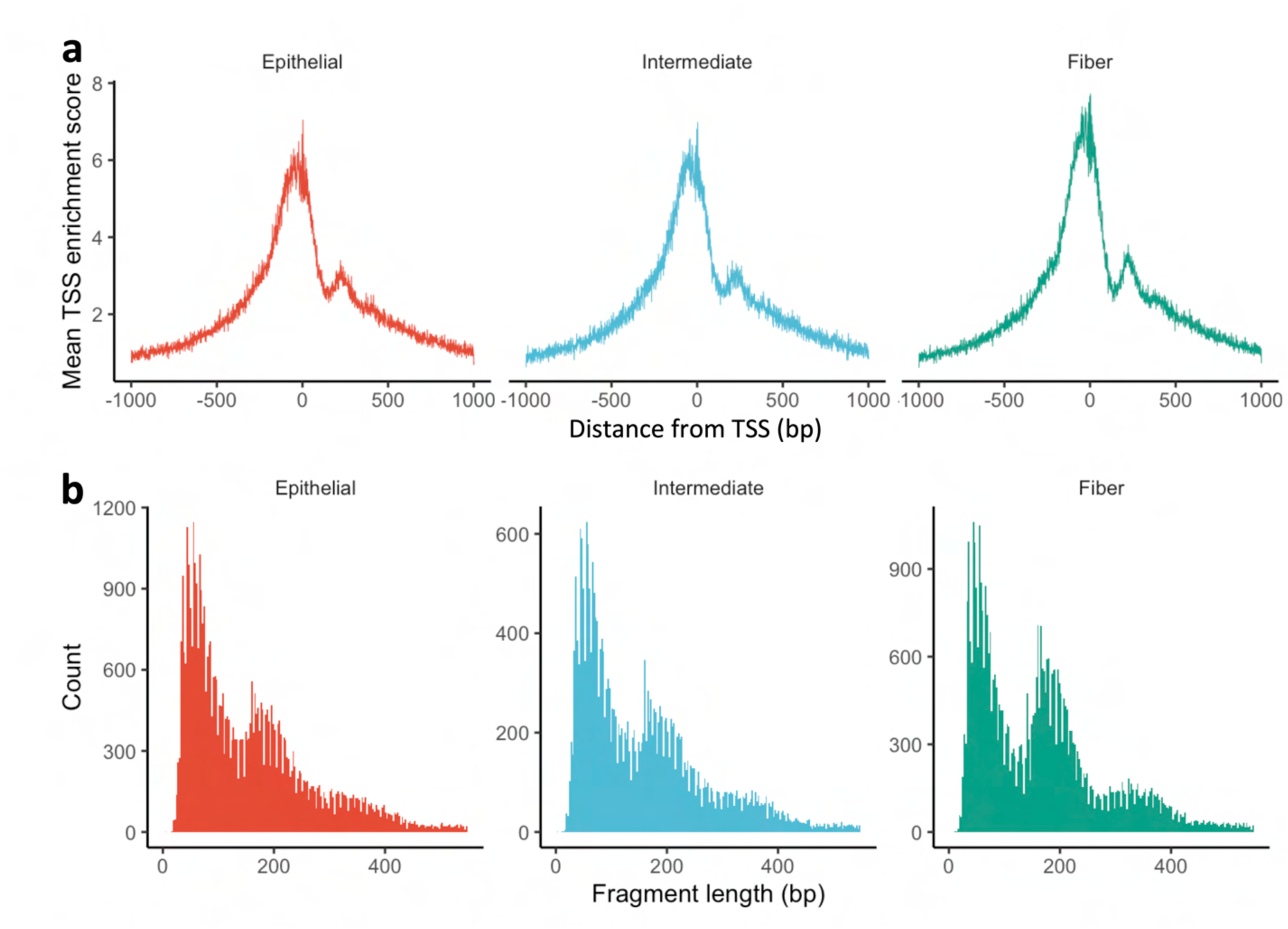
ATAC quality metrics. **(a):** Transcription start site (TSS) enrichment of ATAC signal displayed for each cell population. **(b):** Fragment length of aligned ATAC reads displayed for each cell population. Clear nucleosomal periodicity is observed.

**Supp. fig 3.**
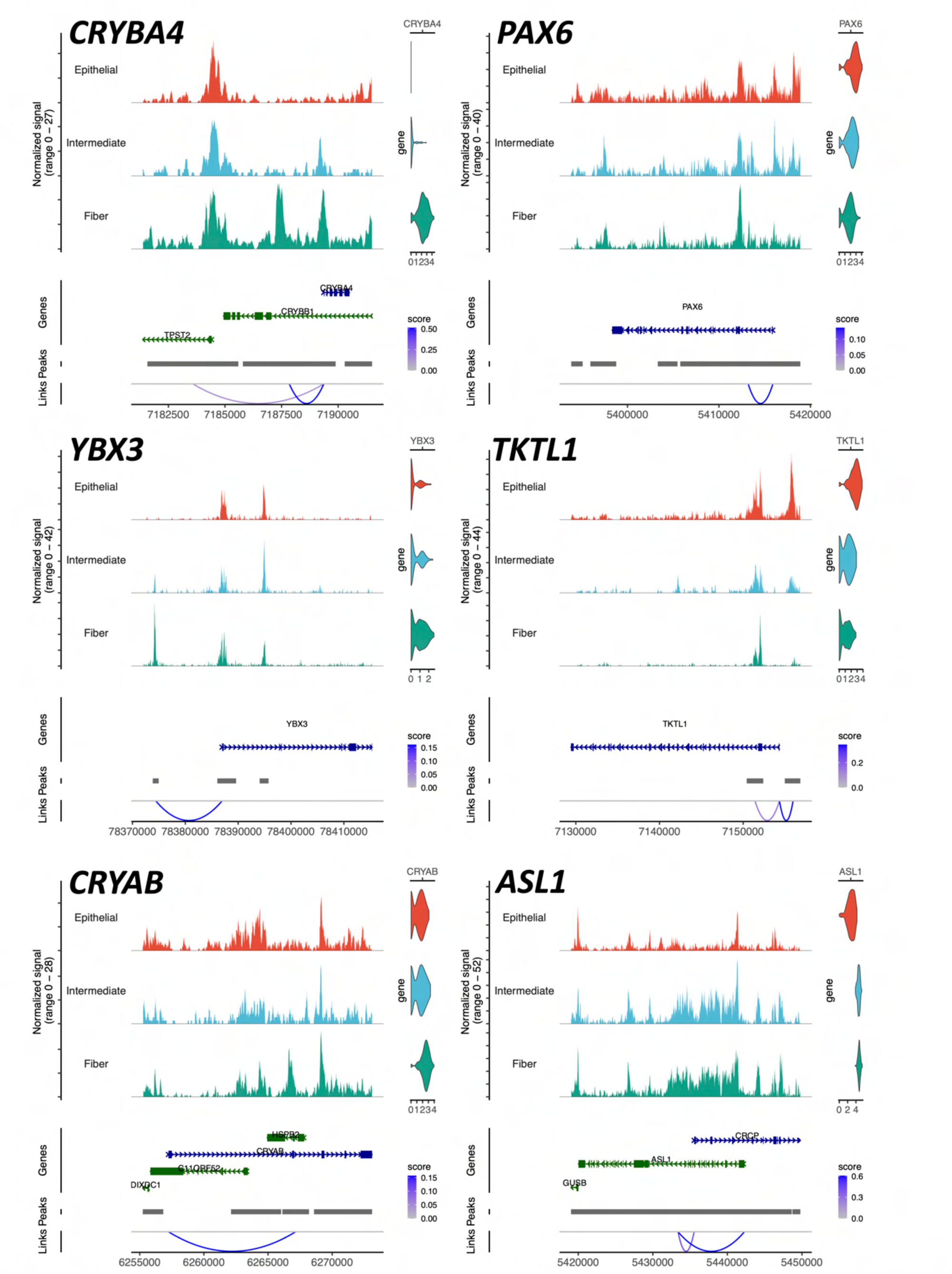
Genome browsers display accessibility signal plotted across genomic loci of interest for each subcluster. The structure of the gene body is indicated by blue bars, with thick regions corresponding to exons, and arrows oriented toward direction of coding sequence. Link tracks on bottom display predicted looping interactions between peak regions and transcription start sites. The links are colored by the link score, which corresponds to the strength of the predicted association between accessibility and expression. Normalized RNA abundance for each gene is displayed in violin plots on right. Numbers along the bottom of each subpanel are genomic coordinates.

**Supp. Fig. 4.**
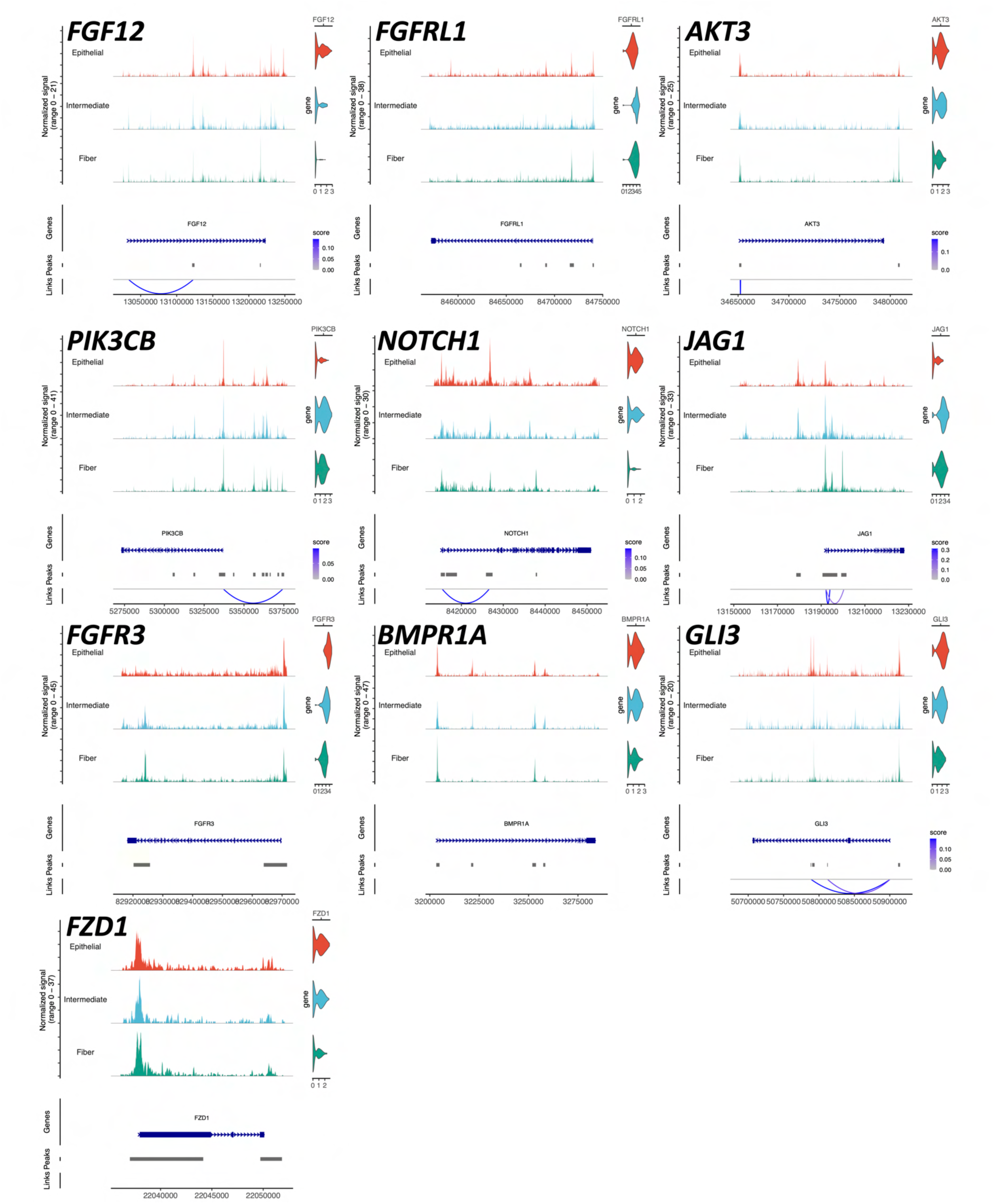
Cis-regulatory activity at signaling effector genes. Genome browsers display accessibility signal plotted across genomic loci of interest for each subcluster. The structure of the gene body is indicated by blue bars, with thick regions corresponding to exons, and arrows oriented toward direction of coding sequence. Link tracks on bottom display predicted looping interactions between peak regions and transcription start sites. The links are colored by the link score, which corresponds to the strength of the predicted association between accessibility and expression. Normalized RNA abundance for each gene is displayed in violin plots on right. Numbers along the bottom of each subpanel are genomic coordinates.

**Supp fig. 5.**
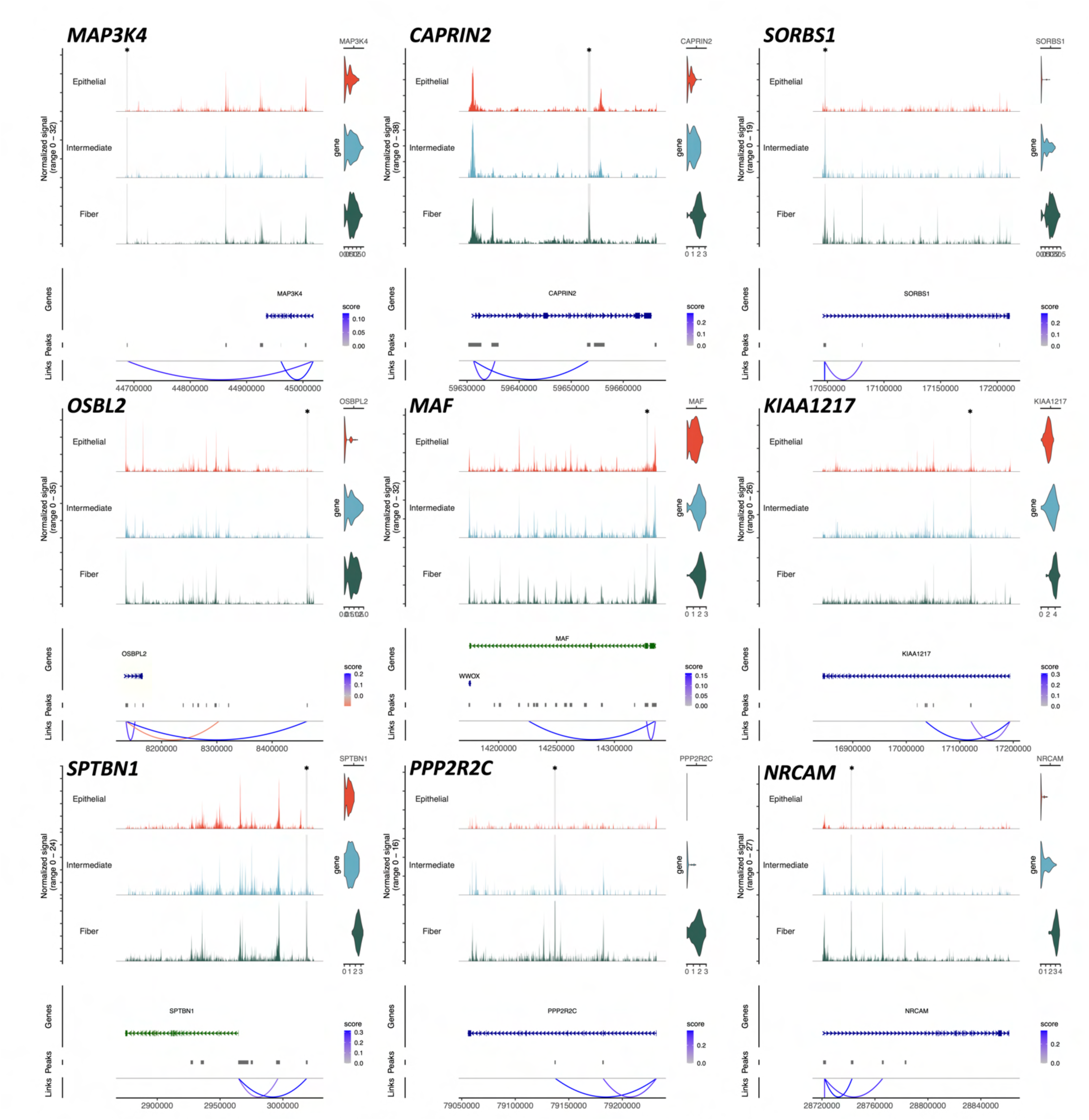
Predicted MAF binding events. The loci of genes containing MAF-linked are displayed, with MAF motif-containing peaks highlighted and marked by an asterisk. Accessibility signal is plotted across the loci. The structure of the gene body is indicated by blue bars, with thick regions corresponding to exons, and arrows oriented toward direction of coding sequence. Link tracks on bottom display predicted looping interactions between peak regions and transcription start sites. The links are colored by the link score, which corresponds to the strength of the predicted association between accessibility and expression. Red links correspond to an inverse relationship between accessibility and expression. Normalized RNA abundance for each gene is displayed in violin plots on right.

**Supp. Fig 6.**
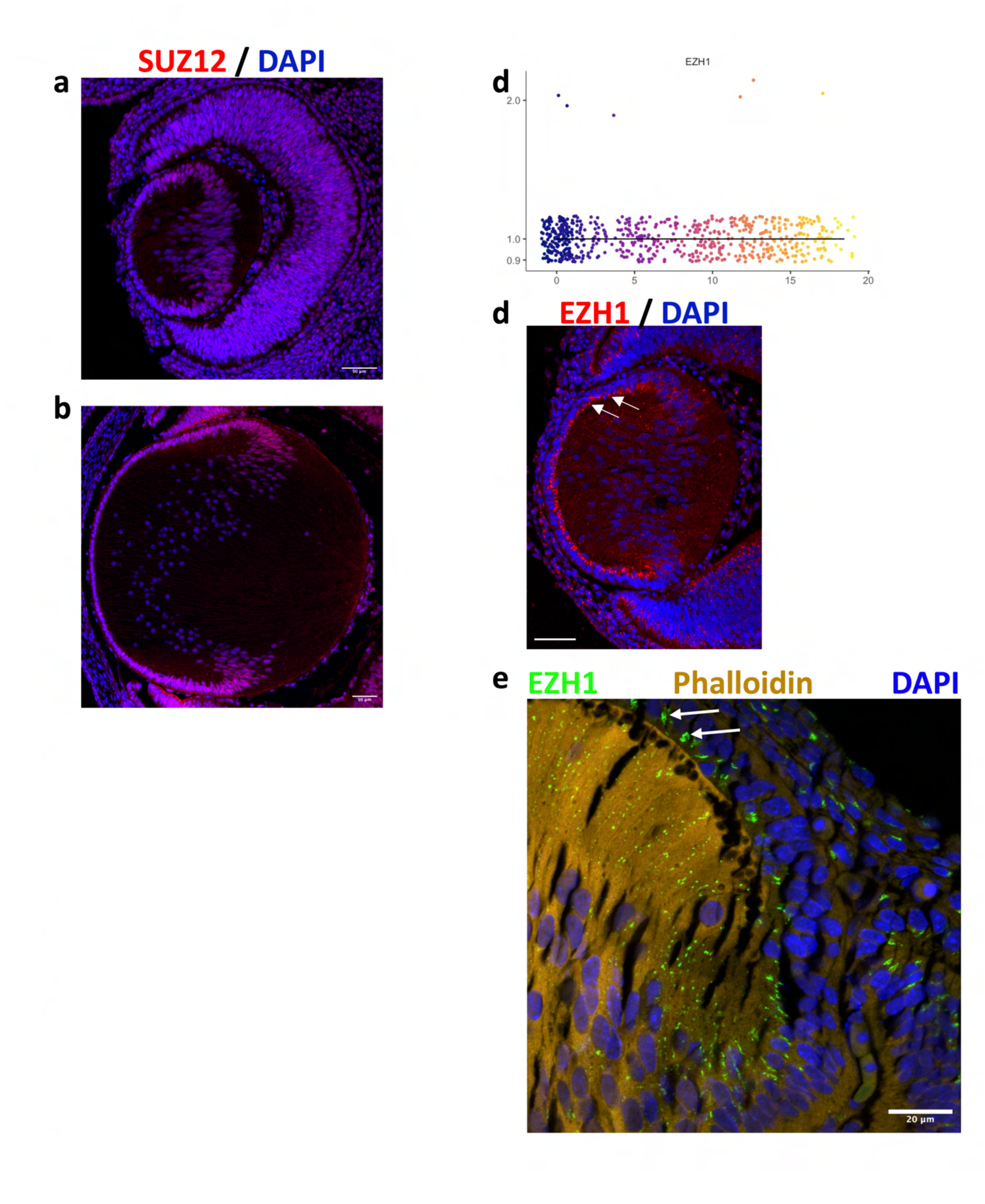
**(a):** An additional replicate of an El2.5 mouse eye stained via IHC against SUZ12, as in Fig. 7C. Scale bar is 50 µm. **(b):** An E14 mouse lens stained via IHC against SUZ12. Scale bar is SO µm. **(c):** *EZHl* is observed to be low-expressed in the snRNAseq dataset and did not display significant evidence of regulation across the epithelial to fiber cell trajectory. **(d):** lmmuno against EZHl using an E12.S mouse lens. Nuclear EZHl staining was not observed. Arrows point to EZHl+ foci observed along the apical surface of lens epithelium. Scale bar is SO µm. **(e):** High-mag image of an El2.5 mouse lens shows EZHl+ foci along epithelial apical surface, indicated by arrows. Phalloidin staining shows F-actin localization. EZHl puncta persist in differentiating fiber cells but disaggregate and redistribute. Nuclei stained with DAPI. Scale bar is 20 µm.

**Supp. Fig 7.**
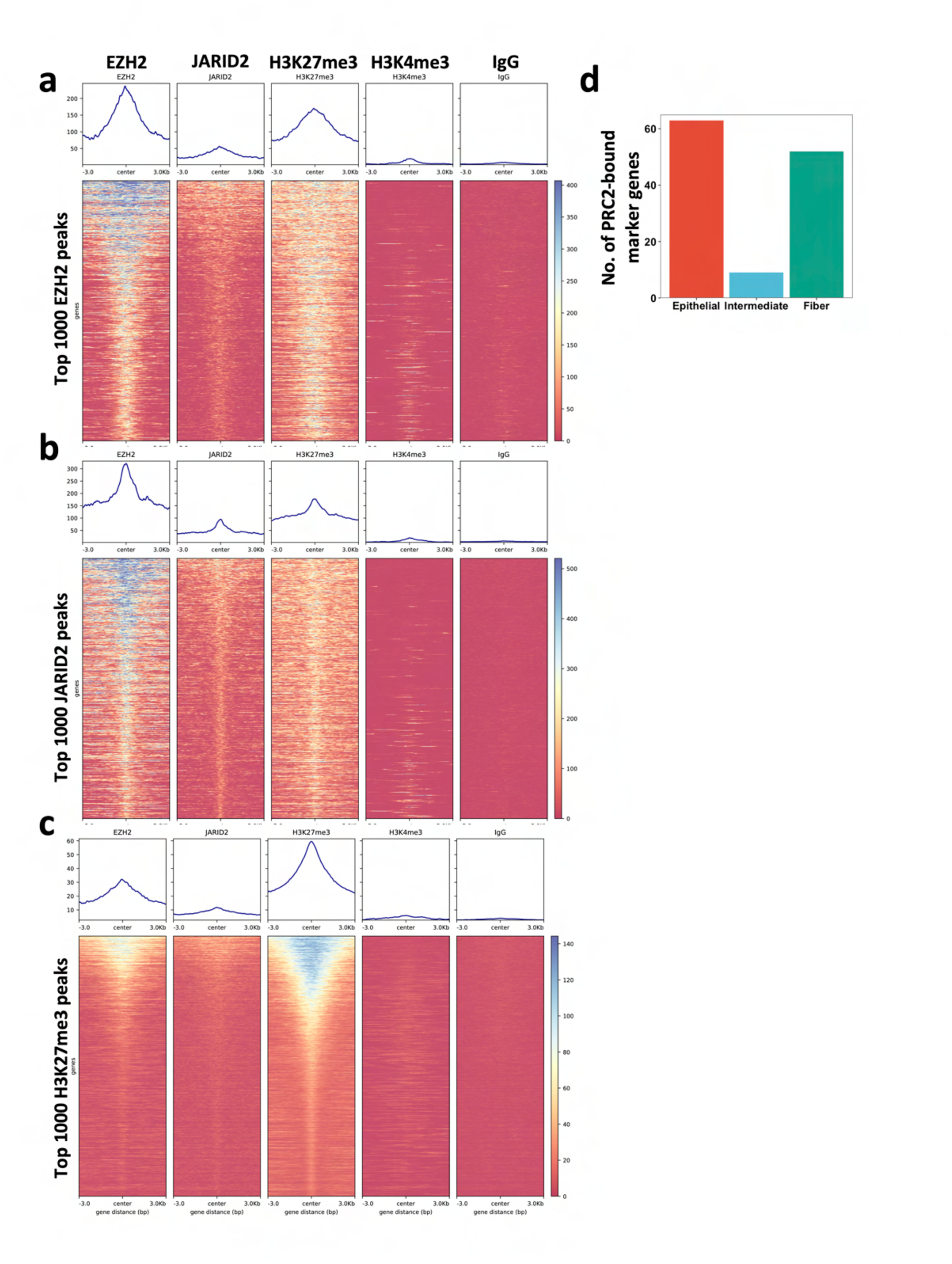
Heatmaps display the observed RPGC­ normalized CUT&RUN-seq signal for indicated marks at specified loci. Loci in **a** are top 1000 EZH2 peaks, **b** top 1000 JARID2 peaks, and **c** top 1000 H3K27me3 peaks. **(d):** Bar chart displays the number of marker genes for each cell population (determined via snRNA-seq) which contain an overlapping JARID2 and EZH2 peak.

**Supp. Fig 8.**
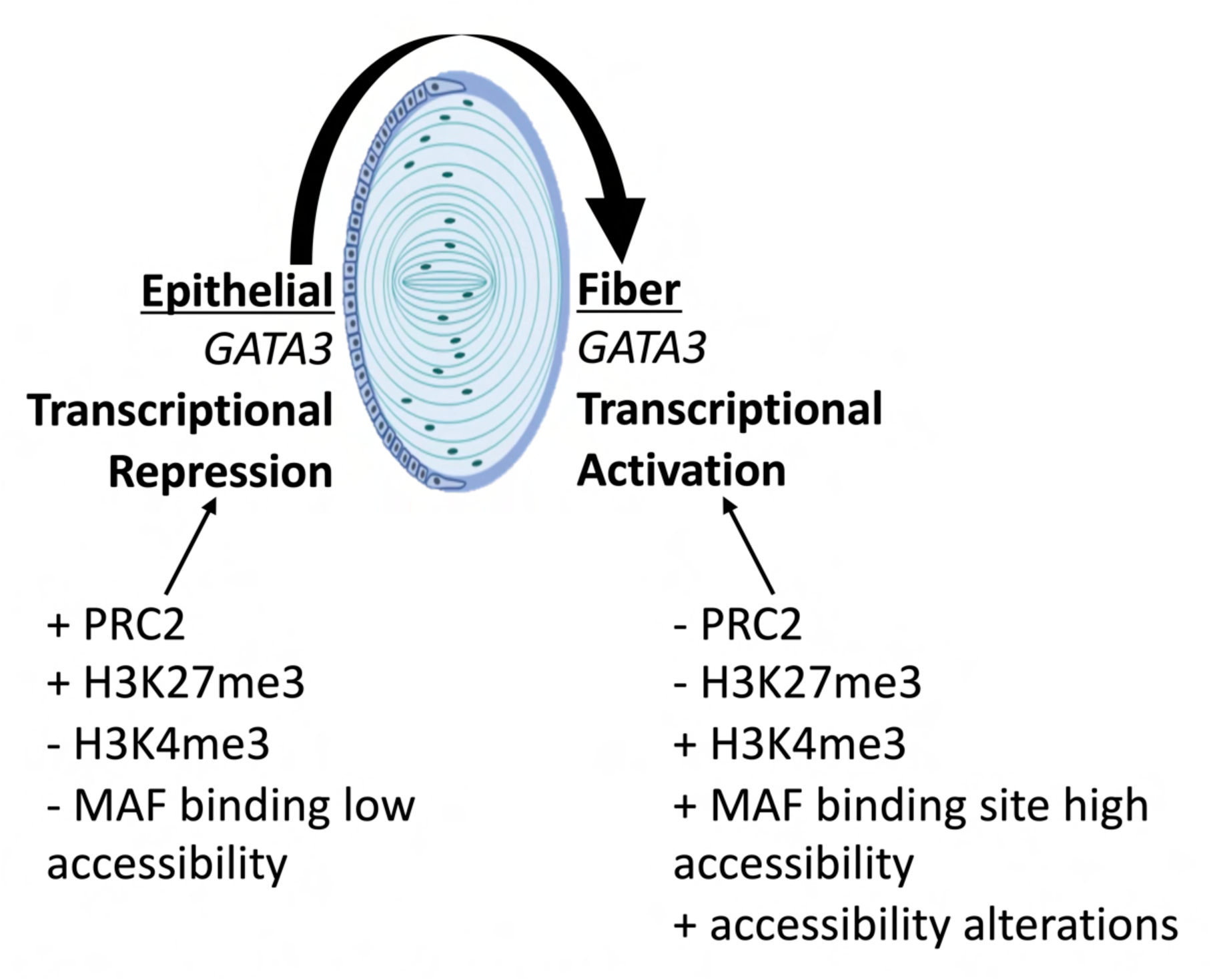
Schematic displays a proposed mechanism for *GATA3* activation in differentiating fiber cells.

**Supp. Fig. 9.**
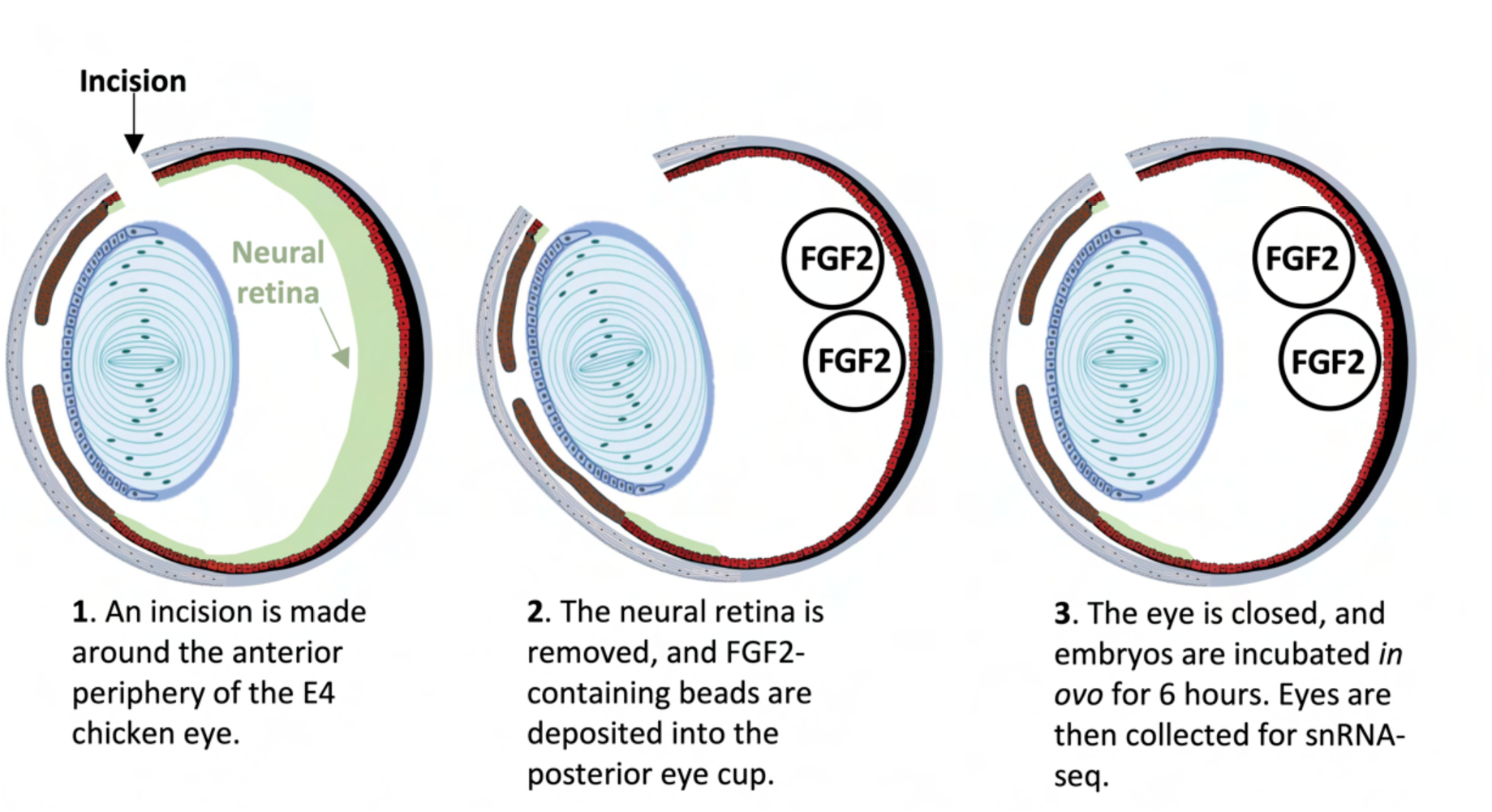
Schematic outlines the surgery performed for *in vivo* delivery of FGF2 to the lens.

**Supp. Fig 10.**
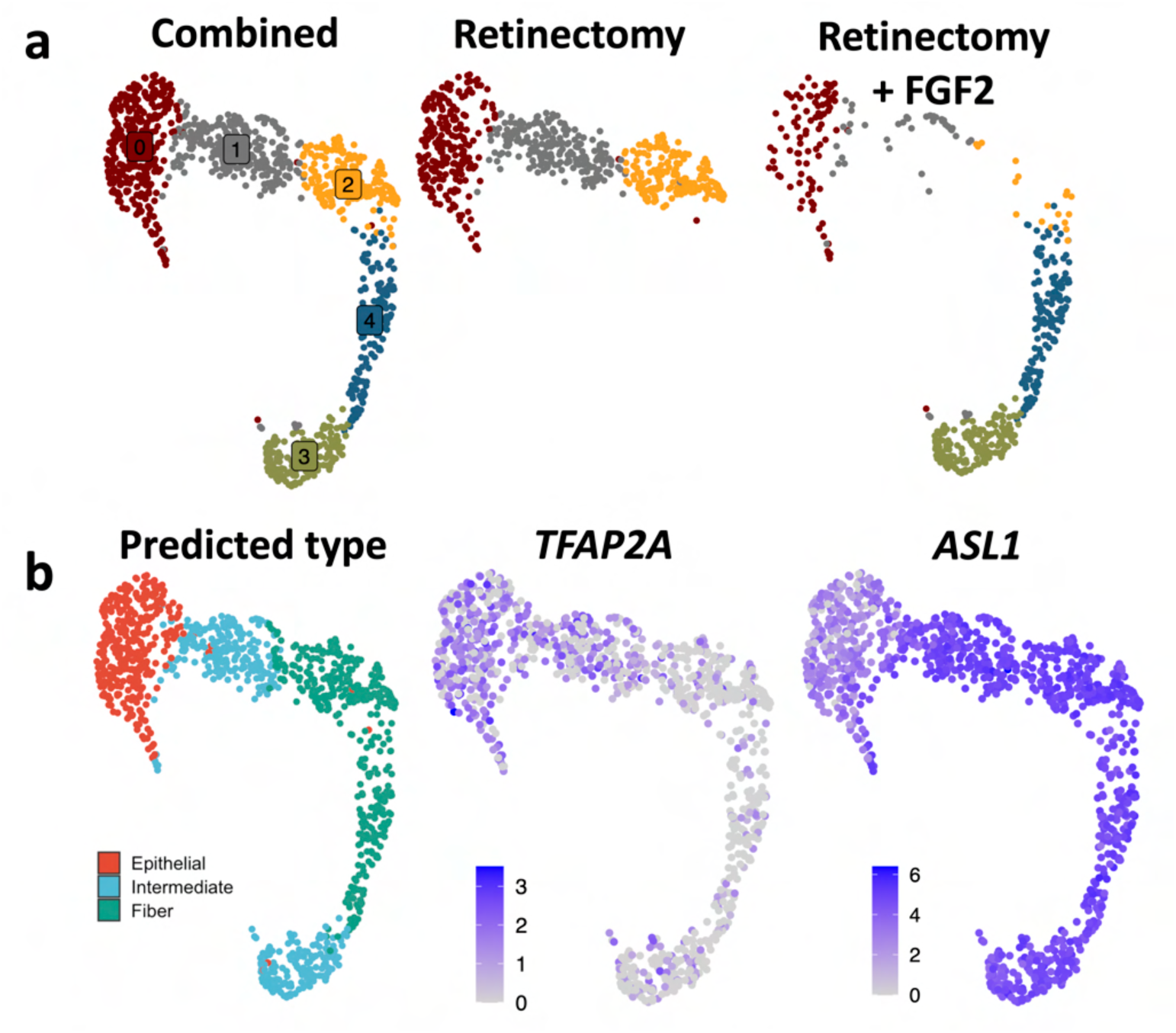
Analysis of the FGF2-treated dataset. **(a):** UMAPs are colored by cluster (clusters numbered 0-4) for all cells (left), retinectomy only (middle), or retinectomy + FGF2 (right). **(b):** UMAPs summarize gene expression across the clusters. Left panel shows cells colored by predicted cell state, middle feature plot is color coded by log-normalized abundance of epithelial gene *TFAP2A,* and right is color coded similarly by abundance of fiber cell gene *ASLl*.

**Supp. Fig 11.**
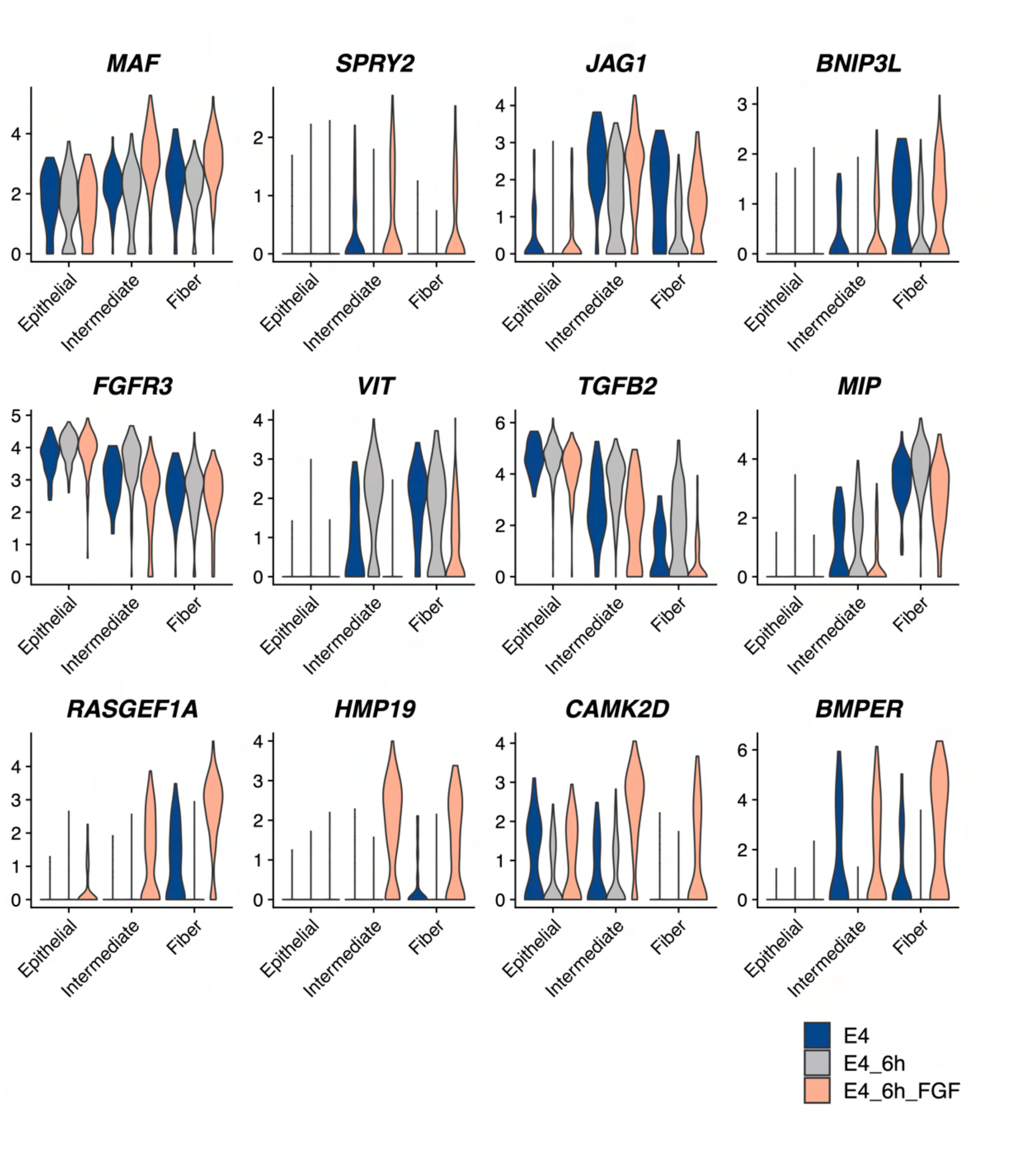
Analysis of the FGF2-treated dataset. Violin plots display the distribution of log-normalized transcript abundance for DEGs across E4 development (Intact), 6 hours post­ retinectomy, and 6 hours post-retinectomy + FGF2 lens populations.

**Supp. Fig 12.**
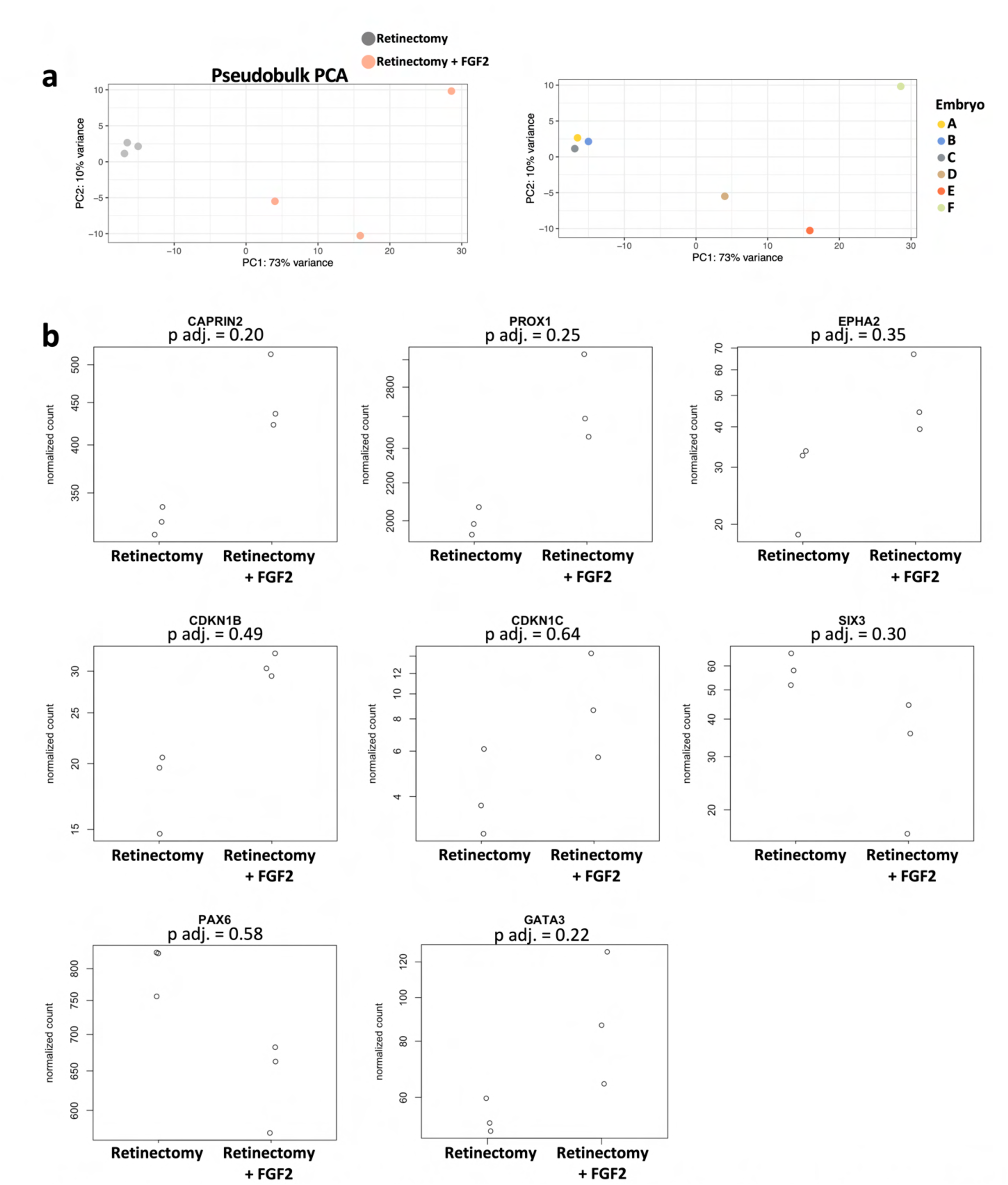
Pseudobulk analysis of FGF2-treated dataset. **(a):** PCA summarizes variation present in lens pseudobulk datasets. In left panel, each biological replicate (embryo) is colored by condition, with conditions separated along PCl. In right panel, points are colored by biological replicate (embryo). **(b):** The displayed genes represent trends in RNA abundance between the conditions which did not achieve statistical significance (adjusted p value> 0.05).

**Supp. Fig 13.**
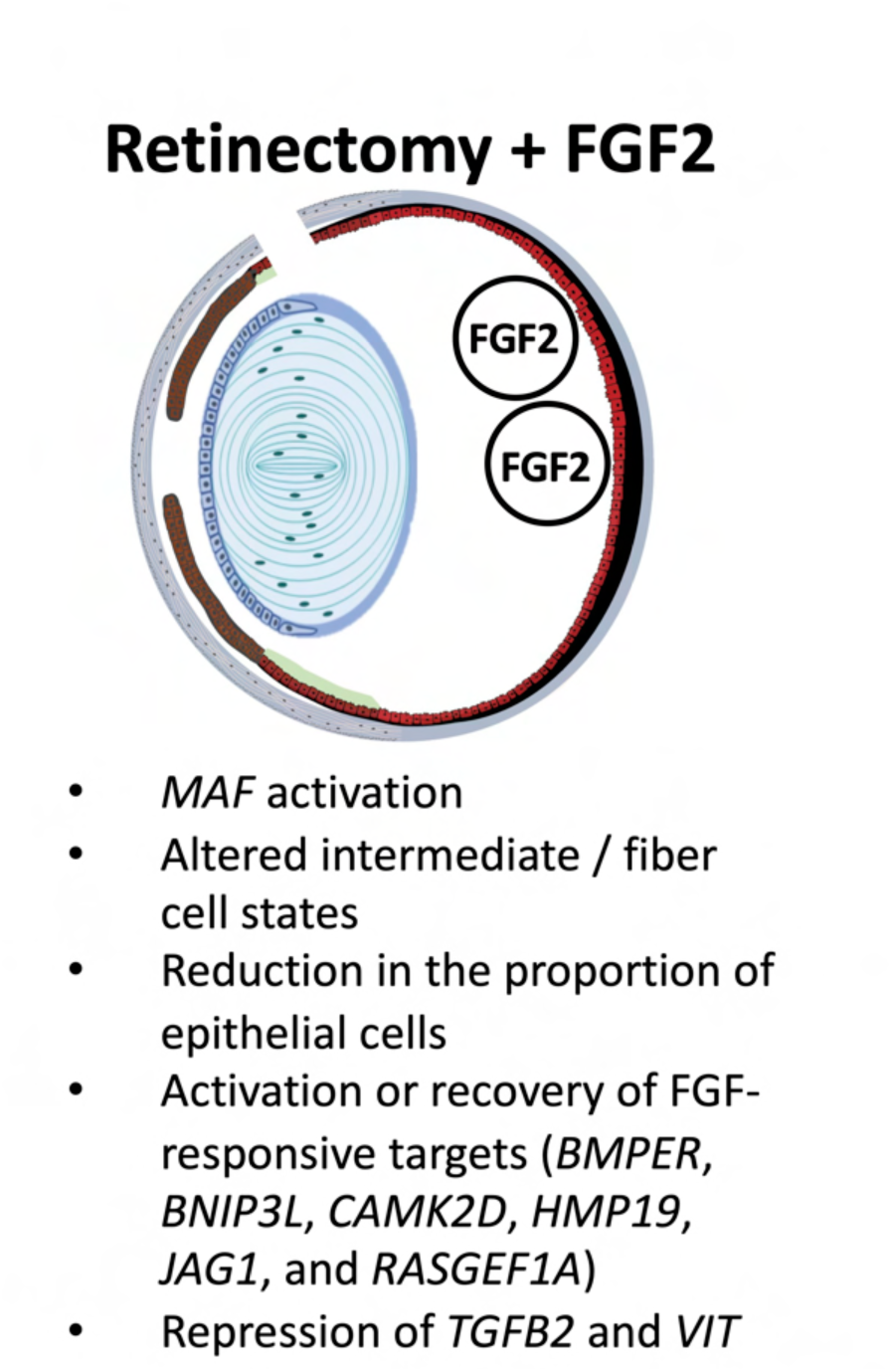
Schematic summarizes observed changes in response to retinectomy and FGF2 treatment.

**Supp. Fig 14.**
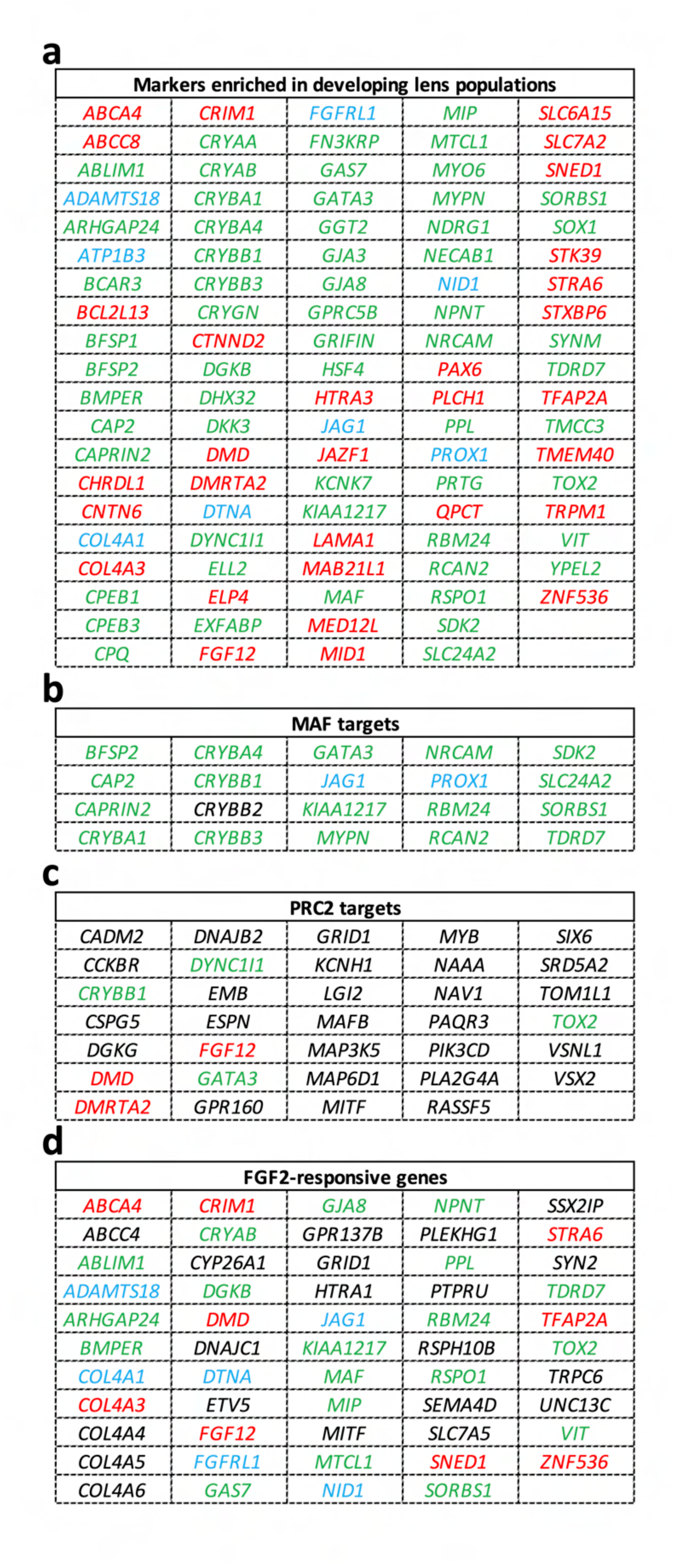
Overlap of genes identified in study and iSyTE. Tables summarize the genes identified in the current study that are documented in iSyTE. Lists encompass significant marker genes drawn from the developing chicken snRNAseq dataset **(a),** genes with regulatory association with nearby predicted MAF binding sites **(b),** genes bound by PRC2 in the developing chick lens **(c),** and genes with significant regulation in response to FGF2 hyperstimulation **(d).** Genes are colored according to their enrichment patterns during lens development: Red = Epithelial­ enriched; Blue= Intermediate­ enriched; Green= Fiber-enriched; **Black= Not significantly enriched in a cluster during development, or not expressed.**

**Supp. Fig 15.**
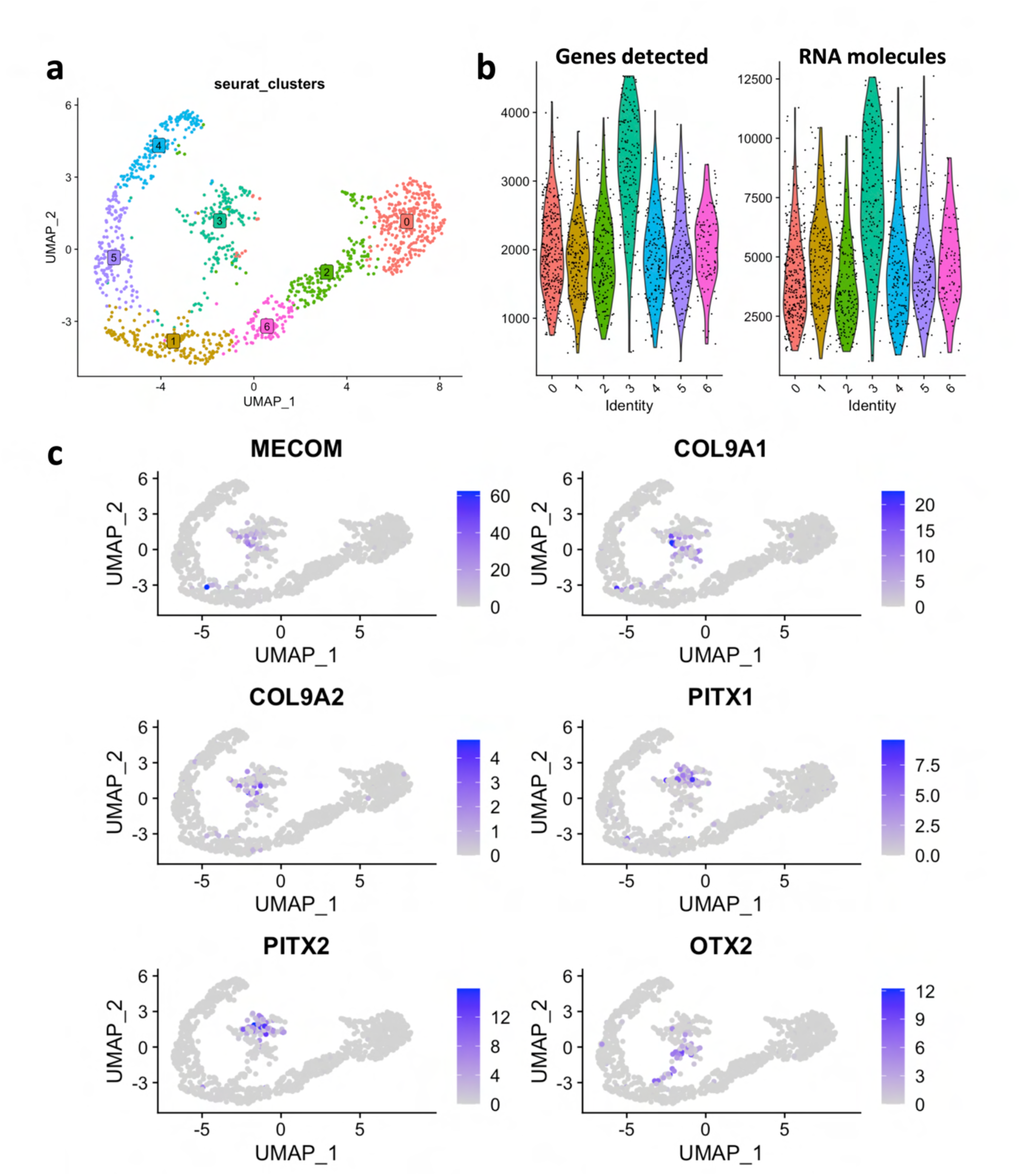
A multiplet population was filtered from the FGF2-treated dataset. **(a):** Lens cells were analyzed as described for the developmental dataset. This process captured a suspected multiplet population (cluster 3). **(b):** The suspect population displayed an aberrantly high number of genes detected and number of RNA molecules per nucleus, characteristic of a multiplet population. **(c):** The suspected population displayed high RNA abundance of cell type-specific genes, including ciliary margin markers *(MECOM, COL9A1,* and *COL9A2),* mesenchyme markers *(PITXl, PITX2),* and RPE / neural retina markers *(OTX2).* The population was excluded from subsequent analysis.

